# A tale of two shrimps – Speciation and demography of two sympatric shrimp species from hydrothermal vents

**DOI:** 10.1101/2024.12.18.629080

**Authors:** Pierre Methou, Shannon B. Johnson, John Sherrin, Timothy M. Shank, Chong Chen, Verena Tunnicliffe

## Abstract

Hydrothermal vents can serve as natural laboratories to study speciation processes due to their fragmented distribution often with geographic barriers between habitats. Two sympatric species of Rimicaris shrimps occur at vents on the Izu-Bonin-Mariana volcanic arc: Rimicaris loihi also occurs near Hawai’i and R. cambonae is present on the Tonga Arc. These two species biogeographically co-occur and are genetically similar, raising questions about their speciation mechanism, how they maintain distinct species, and whether interbreeding occurs. Here, we use barcoding and shotgun sequencing to test their genetic isolation and investigate their speciation process. We also evaluate population demography over 10 years to assess population densities and sex ratios at vents. Our results support R. cambonae and R. loihi as two distinct species despite sympatry throughout part of their range. We also observed regional-scale genetic structure among R. loihi populations from the Izu-Bonin-Mariana volcanic arc despite their broad geographic distribution. Finally, we found concomitant variations of shrimp densities and genetic diversity following fluctuations in geological and venting activities over a decade. A combination of geological instability, ocean currents dynamics and sea-level changes might drive temporary isolation among these local populations. We suggest similar factors, with longer isolation periods, may also have promoted speciation between the two Rimicaris species whereas distinct life-history traits could strengthen and maintain reproductive barriers. Overall, our results indicate different connectivity patterns on a volcanic arc that contrast with those observed previously at vents from mid-ocean ridges or back-arc basins systems.

## 1. Introduction

Hydrothermal vent habitats on ocean ridges and volcanic arcs share characteristics with open ocean island archipelagos: they predominantly host endemic species, are isolated by expanses of inhospitable seafloor, and vulnerable to shifts in local conditions and the arrival of colonizing species. The dynamics of speciation on ocean island systems distal from mainland effects have been substantially explored in theoretical ecology (MacArthur & Wilson, 2001; Whittaker et al., 2008) amplified by studies on temporal processes and on regional phenomena such as volcanism or sea-level fluctuations (Fernández-Palacios, 2016). Much less is known about marine island-like settings. Seamount chains rising from the abyss, sometimes to sunlit depths, are possible analogies as the depth differences between them acting as dispersal barriers. Compared to the surrounding ocean floor, the upper slopes and crests of seamounts are often diversity hotspots (Morato et al., 2010). For the most part, seamount species are broadly distributed regionally although populations of some species show relatively poor connectivity; endemics are rare (Rogers, 2018). Marine organisms tend to have higher dispersal capabilities compared to their terrestrial counterparts, due to swimming adults or pelagic larval stages. Consequently, new subsea “islands” accumulate new species mostly by migration. Poor dispersers can speciate in isolated localities when gene flow diminishes, but radiation is deterred by new colonizers that may out-compete or interbreed with the incipient species (Pinheiro et al., 2017).

Hot vent ‘archipelagos’ along oceanic ridges, volcanic arcs, and back arc basins are notable for their high endemism (Tunnicliffe et al., 1998). Within a vent field, the steep thermal and chemical gradients generated by the mixing of vent fluids with seawater create a variable mosaic of distinct habitats constraining the spatial distribution of faunal communities (Cuvelier et al., 2011; Podowski et al., 2010). Even after45 years of both vent site and species discovery, rates of reported endemism have not diminished (Beaulieu & Szafrański, 2020; Tunnicliffe et al., 2023). Across eleven vent systems (= “archipelagos”) of the western Pacific, two-thirds of the species occur in only one system despite, in some cases, their close proximity; frequently, species in the same genus occupy identical niches (Tunnicliffe et al., 2023). While connectivity studies at vents focus on taxa with broad ranges along mid-ocean ridges or back-arc basins, we know little about the speciation dynamics of poor dispersers or the outcomes of secondary contact by long-distance dispersers (Breusing et al., 2020; Castel et al., 2022; Johnson et al., 2013; Matabos & Jollivet, 2019; Thomas-Bulle et al., 2022). Connectivity of species living only at vents on volcanic arcs have also received less attention. Overall, the biogeographic histories and responses of marine organisms living in underwater “islands” may differ significantly from terrestrial counterparts living on islands. Case studies from hot vents, as key examples of underwater archipelagos, provide the basis for comparative assessments.

Among the vent-endemic fauna, species of Rimicaris shrimps are commonly found living in sympatry in several regions, where two or three species can co-occur within the same vent fields (Lunina & Vereshchaka, 2014; Methou, Hernández-Ávila, et al., 2022; Methou, Chen, et al., 2024). In most cases, Rimicaris shrimps exhibit broad geographic distributions with high genetic connectivity along mid-ocean ridges (Beedessee et al., 2013; Teixeira et al., 2012; Zhou et al., 2022) or back-arc basins (Methou, Ogawa, et al., 2024; Thaler et al., 2014; Yahagi et al., 2015), suggesting a high dispersal potential. As with most decapods, Rimicaris shrimps are gonochoric brooders that maintain their eggs under their abdomen but also display a variety of reproductive cycles with seasonal or aperiodic brooding patterns (Copley & Young, 2006; Methou, Chen, et al., 2022; Methou et al., 2023). In some Rimicaris species, brooding cycles were unrelated to variations in photosynthetic surface production, unlike other seasonal brooders that inhabit vents (Methou, Chen, et al., 2022).

Two sympatric Rimicaris species co-occur at vents from the Izu-Bonin-Mariana (IBM) Volcanic Arc in the Northwest Pacific: Rimicaris (formerly Opaepele) loihi and the recently described R. cambonae (Methou, Chen, et al., 2024). The hydrothermally-active habitats of R. loihi and R. cambonae are highly diverse and volatile, characterized by areas of liquid CO_2_ emergence at NW Eifuku, molten sulfur ponds at Nikko Seamount and other sites with an extensive record of historical eruptive events (Embley et al., 2007; Lupton et al., 2006). In particular, active eruptions occurred at NW Rota Volcano during each of the six visits between 2004 and 2010 (Chadwick et al., 2008, 2012; Embley et al., 2006; Schnur et al., 2017). Rimicaris loihi was also reported from distant sites including Loihi Seamount off the southeast coast of Hawai’i (Williams & Dobbs, 1995) and at West Mata Volcano north of Tonga (Resing et al., 2011). West Mata and Loihi are also unstable due to seismic and eruptive activities, as recorded at Loihi in 1996 and by in situ observations of explosive eruptions during dives at West Mata in 2008 and 2009 (Garcia et al., 2006; Resing et al., 2011).

The sympatric R. cambonae and R. loihi are also genetically similar, differing by just 2% in the barcoding region of the mitochondrial cytochrome c oxidase subunit I (COI) gene fragment (Methou, Chen, et al., 2024) and thus falling at the broad border between divergent populations and species (Hebert et al., 2003). Given that they coexist within the same faunal assemblages, questions arise about the presence of gene flow and, thus, the effectiveness of reproductive barriers despite the potential for reproduction. However, data on the reproductive biology of these species are limited to one single population of R. loihi from Suiyo collected in August 2019, that revealed a largely female-biased sex ratio and presence of many juveniles and ovigerous females (Methou et al., 2023).

This study aims to answer the following questions: 1) Are nuclear genes of R. loihi and R. cambonae concordant with mitochondria and represent separate species-level lineages?; 2) Is there gene flow and hybridisation between R. cambonae and R. loihi or is there a complete reproductive isolation?; 3) To what extent does the eruptive and unstable geological context affect the population dynamics of these shrimps over the years?; and 4) What is the most likely scenario for their speciation?

## 2. Materials and Methods

### 2.1 Animal sampling and processing

Alvinocaridid shrimps were collected during eight oceanographic expeditions between 2004 and 2023 using suction samplers or baited traps on either Remotely Operated Vehicles (ROVs) or Human Operated Vehicles (HOVs). In total, five hydrothermal vent fields from the Izu-Bonin-Mariana (IBM) Volcanic Arc were visited once (Seamount X in 2006, Suiyo in 2019), twice (Nikko in 2006 and 2010, NW Eifuku in 2004 and 2023) or six times (NW Rota between 2004 and 2014) (Figure 1A). Additional shrimps were collected in 2004 at Loihi Seamount near Hawai’i, the type locality of Rimicaris loihi and in 2009 at the West Mata volcano (Figure 1A; see Table S1 for detailed sampling information).

**Figure 1.**
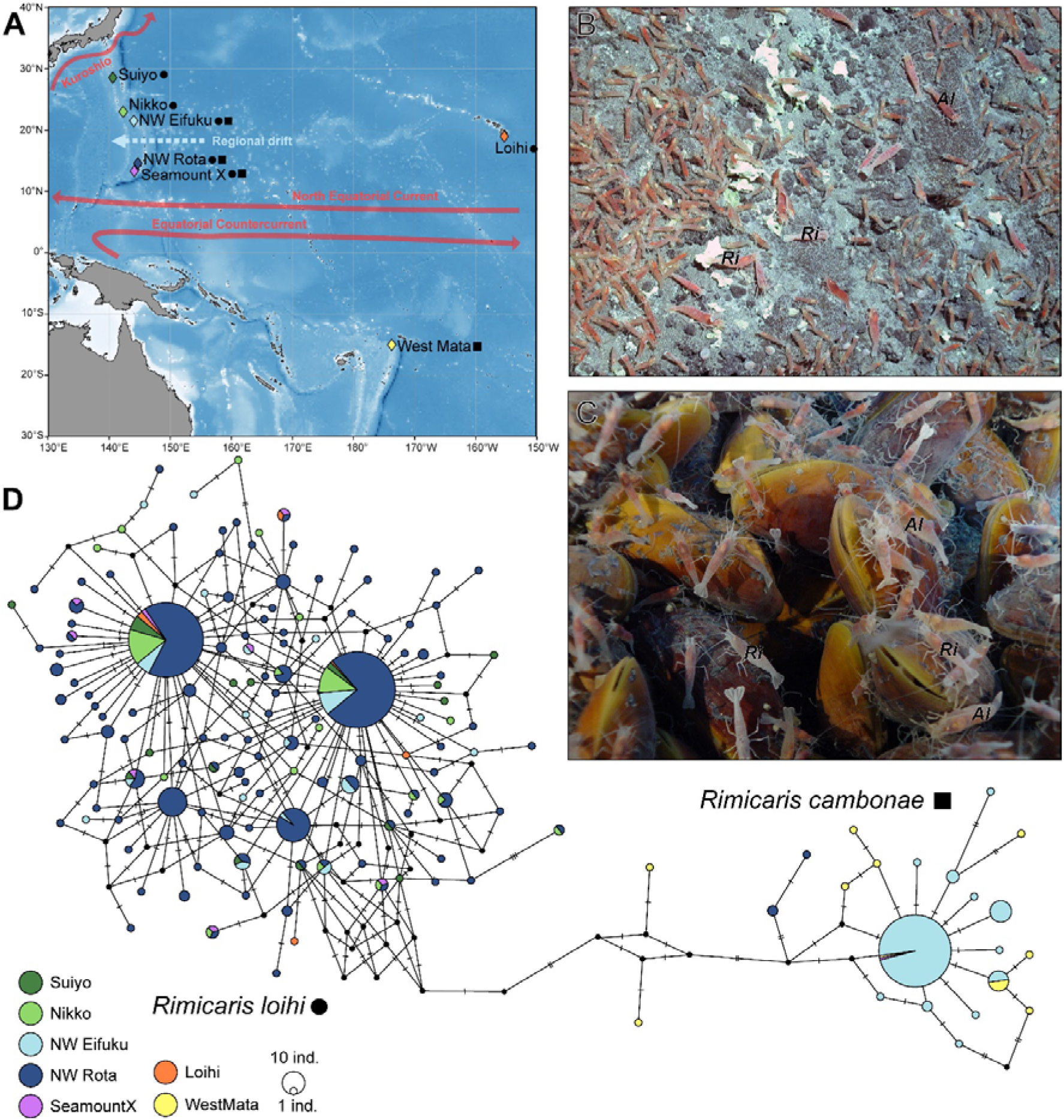
**A.** Geographic distribution of samples for Rimicaris cambonae (square) and Rimicaris loihi (circle). Coloured diamonds indicates localities and black shapes (circles and squares) indicates species. Typical habitats of Rimicaris spp. (Ri) on **(B)** sulfur crusts (NW Rota, 2009) and **(C)** mussel beds (NW Eifuku, March 2004). **D.** COI haplotype network (529 bp) of Rimicaris species from Loihi, West Mata and the Mariana Arc vent fields (n= 597) detailed per sampling site. Two clusters emerge with 24 haplotypes affiliated with R. cambonae and 134 haplotypes affiliated with R. loihi.

Specimens were preserved either in 75% ethanol or frozen whole at -80°C on-board the ship. Additional specimens from NW Eifuku in 2023 were fixed with 4% paraformaldehyde (PFA) for 3h on board and stored in 50% phosphate-buffered saline (PBS)-ethanol solution for electron microscopy (see Supplementary Methods). We determined sex based on examination of 1^st^ and 2^nd^ pleopod shape and measured the size of each shrimp as carapace length (CL) and/or total length (TL) following (Methou, Chen, et al., 2024). For the few ovigerous females encountered, we estimated realized fecundity from egg counts and sizes. For 144 specimens collected in 2023 at NW Eifuku, we distinguished R. cambonae and R. loihi based on morphological characters defined by (Methou, Chen, et al., 2024) – briefly, the form of the pterygostomial tooth on the front part of the carapace and the shape of the proximolateral tubercle on the antennular stylocerite – in conjunction with COI barcoding. However, as specimens from earlier work were no longer available for morphological study, species identification reported below relies on genetic assessment.

### 2.2 Shrimp Abundance

On NW Rota, vents occur in the upper 80 m of the summit. Fine-scale mapping with an SM2000 system, good ROV navigation, and physical markers (e.g., Schnur et al. 2017) supported comparative estimates of volcanic activity and venting extent between years. However, rugged terrain and radical changes in the volcano topography hampered repeated transects. Instead, we estimated shrimp abundance each year by assessing the total fraction of the ROV transects in which shrimps were visible in densities greater than ∼20 individuals/m^2^.

### 2.3 DNA extraction and Sanger sequencing

We performed DNA extractions on 597 alvinocaridid shrimps using the 4^th^/5^th^ pleopods or pieces of abdominal tissues following manufacturer’s instructions of the DNeasy Blood & Tissue kit (Qiagen, Hilden, Germany). We amplified the 5’ region of the cytochrome c oxidase subunit I gene (COI) with the Cari-COI-1F and Cari-COI-1R alvinocaridid primers (Methou et al., 2020) for shrimps collected from Nikko 2010, Suiyo 2019, and NW Eifuku 2023 (679 to 734 bp fragment) or with the COI-1f (5’-AGTAGGAACAGCCCTTAGTCTCCTA-3’) and COI-1r (5’-TCCCGCTGGGTCAAAGAATGAGG-3’) primers for the remaining shrimps (529 to 623 bp fragment).

We used Promega GoTaq polymerase in 25µl reactions for PCR amplifications with Cari-COI-1F and Cari-COI-1R and iProof High Fidelity DNA Polymerase (Bio-Rad) in 40µl reactions for PCR amplifications with COI-1f and COI-1r using 1 µl of the DNA template for both. We conducted PCR with the following cycle conditions: initial denaturation at 95°C for 5 min, 35 cycles of 95°C (1 min), 50°C (1 min), 72°C (2 min), followed by a final extension stage of 72°C for 7 min with Cari-COI primers or initial denaturation at 95°C for 5 min, 35 cycles of 95°C (1 min), 50°C (1 min), 72°C (2 min), followed by a final extension stage of 72°C for 7 min with the COI-1f/ COI-1r primers.

Bi-directional Sanger sequencing was conducted by Eurofins (Köln, Germany) for PCR products amplified with Cari-COI-1F/ Cari-COI-1R primers and by the High Throughput Genomics Unit (Seattle, WA, USA) for PCR products amplified with COI-1f/COI-1r primers. We edited and aligned sequence chromatograms with Geneious Prime® 2023.1.2 (https://www.geneious.com).

### 2.4 Haplotype networks

To assign individual mitochondrial barcodes to putative species, we used the Assemble-Species-by-Automatic-Partitioning method on a 529-bp alignment, a method based on the detection of “barcode gaps” (Puillandre et al., 2021). We estimated median joining haplotype networks with PopART v1.7 (Bandelt et al., 1999; Leigh & Bryant, 2015) with default settings. We used DnaSP v6 (Rozas et al., 2017) to determine the number of variable sites (S), the number of haplotypes (h), haplotype diversities (H_d_), nucleotide diversities (π), average number of nucleotide differences (k, (Tajima, 1983), raw genetic distance (D_a_), and pairwise F_ST_ values with 1000 permutations within and among putative species and populations.

### 2.5 Whole Genome Shotgun Sequencing and Analyses

#### 2.5.1 Sequencing

Following COI barcoding, we used whole genome shotgun (WGS) sequencing on a subset of randomly selected animals of each species along our sampling range on the IBM volcanic arc to examine whether nuclear data were concordant with mitochondrial data and to explore connectivity amongst populations and lineages. We quantified genomic DNA with the DNA High-Sensitivity assay with the Qubit 2.0 Fluorometer (ThermoFisher Scientific, Waltham MA, USA) and prepared 32 individuals for WGS sequencing with the Illumina DNA Prep kit (Illumina, Inc. San Diego, CA). We sequenced six randomly chosen individuals of R. loihi from Suiyo and Nikko, eight from Eifuku, and four from NW Rota Seamounts. We also sequenced eight R. cambonae from Eifuku Seamount. Sequences were generated on the Element Bio AVITI instrument at UC Davis with a paired-end 2x150 protocol. Low coverage individuals, including all four specimens from NW Rota were unsuccessful and were excluded from analyses.

#### 2.5.2 Quality filtering

We first removed sequencing barcodes and adapters from WGS data with Trimmomatic v.0.35 (Bolger et al., 2014). To perform population genetic analyses on WGS sequencing, we filtered mitochondrial and PhiX sequencing control data from nuclear reads with the programs BWA MEM v.0.7.17, seqtk, and SAMTools v1.7-1. We then assembled a nuclear genome from the individual that had the highest number of reads (OC_Ebi50, R. cambonae from Eifuku Seamount) with SPAdes (v.3.15.5-Linux, (Prjibelski et al., 2020) with default settings. We examined the assembly with the program BandageNG v.2022.8-dev (https://github.com/asl/BandageNG) and removed contigs <1500 bp long.

We mapped reads from genomic data to the nuclear genome that we assembled for the Ebi50 individual with the program ANGSD v0.940 (Korneliussen et al., 2014). Average read depth was plotted against the sequencing read score quality for all individuals, which enabled us to visualize the data and limit low quality reads, paralogs, and repetitive regions of the genome. Low-coverage individuals, including one R. loihi and R. cambonae each from NW Eifuku and all four specimens from NW Rota were excluded from analyses. We took a number of steps to filter our data within ANGSD to only include high-quality reads. We discarded reads that had multiple hits to the reference assembly (paralogs) and limited read depths from 10-150x coverage where we discarded reads <10X and >150x coverage. We increased SNP site mapping quality by filtering excessive mismatches (C=50), removed all sites with missing data, excluded improperly paired reads, and adjusted qscores around indels to resolve false variants. Each population was individually tested for deviations from the Hardy-Weinberg Equilibrium (HWE, p-value <0.05) and sites that were statistically significant were excluded from subsequent analyses.

#### 2.5.3 Statistics and demographic analyses

We used the programs ANGSD, PCAngsd (Meisner & Albrechtsen, 2018), ngsTools v3.0 (https://github.com/mfumagalli/ngsTools), ∂a∂i v. 2.3.7 (Gutenkunst et al., 2009), StAMPP (Pembleton et al., 2013) and dadi-pipeline v.3.1.7 (Portik et al., 2017) to estimate population and species differentiation, admixture, and demographic histories of populations. With ANGSD, we estimated genotype likelihoods and the minor allele frequency (MAF) to calculate the site frequency spectrum (SFS), to estimate population differentiation; (F_ST_, Reynolds et al., 1983), and summary statistics with at least five individuals per site. Lacking an ancestral sequence to polarize data, we calculated Tajima’s D, Waterson’s Θ, and Nucleotide diversity (π) from the SFS with a ‘folded’ spectrum where each SNP was present in at least five individuals and had a minimum minor allele frequency of 0.05. We also used the R package StAMPP (v.1.6.3, (Pembleton et al., 2013) to calculate F_ST_ values, 95% confidence limits, and p-values among populations and species. To do so we estimated the SFS within ANGSD for the entire dataset with the same filtering parameters as above for sites that at least ten individuals had data and with the -doSaf (1) flag which calculates the site allele frequency likelihood based on individual genotype likelihoods assuming HWE, -doPost (1) flag which uses the allele frequencies as a prior, the -doMaf (1) flag which calculates fixed major and minor per-site frequencies, the -doMajorMinor (1) flag which infers major and minor allele frequencies from the genotype likelihoods, the -doGeno flag (32) which writes the posterior probabilities of the genotypes as binary with a snp_pvalue of 1e^-6^. We then used realSFS to estimate the site frequency spectra, then imported these within a second run of ANGSD, with the same parameters but also including the -doPest flag with the results from realSFS and the -doBcf flag which exports a BCF file. We then used BCFtools (v.1.11-21-g0987715, (Danecek et al., 2021) to convert the bcf file to a vcf file for import to R. Within R, we used vcfR (v.1.15.0, (Knaus & Grünwald, 2017) to import the VCF file for analyses in StAMPP.

We used the dadi_pipeline to infer the joint demographic histories of populations with the folded joint allele frequency spectrum estimated from ANGSD and realSFS from each population SFS estimate. To create an input file for ∂a∂i we formatted the outfile with the perl script: realsfs2dadi.pl (https://github.com/z0on/2bRAD_denovo/blob/master/realsfs2dadi.pl). We tested the three-population pipeline but analyses never concluded, so instead we used the dadi-pipeline two-population script in a pairwise fashion where the spectra were folded (Charles et al., 2018; Portik et al., 2017). First, we compared the sympatric populations of R. loihi and R. cambonae from Eifuku seamount. Second, we compared the allopatric populations of R. cambonae from Eifuku and R. loihi from Nikko seamount. We chose the Nikko population because it was geographically closest to the Eifuku locality. Third, we compared each R. loihi population. We used the dadi-test-projections.py script from the dadi_pipeline to determine the number of projections to maximize the number of segregating sites. We tested both the ‘diversification model set’ (Portik et al., 2017) and ‘island model set’ (Charles et al., 2018) for two populations. The diversification set tests models that range from no migration to more complicated models that include secondary contact and population size changes. We also included the island model diversification set since hydrothermal vents are often compared to isolated island habitats. Dadi_pipeline analyses were run with the default settings from 10-40 replicates for four optimization rounds each, where at the end of each round, the parameters of the best-scoring replicate are used to generate the starting parameters of the next round. Once completed, the lowest AIC value was compared with the Summarize_outputs.py script to find the model that best-fit our data. We plotted results with the Make_plots.py script.

We estimated principal component analysis and population admixture of individual genotypes probabilities with ANGSD with the same filtering parameters as above but also removed sites that were not in HWE. Within ANGSD, we used a MAF of 0.05 and sites where at least 5 individuals had SNP data with a p-value of 10^-3^ to estimate a covariance matrix for PCA for PCAngsd and NGSadmix v.32 (Skotte et al., 2013). We ran NGSadmix five times for each population (K) value and then used the method of Evanno et al. (2005) to compute the most probable number of clusters for NGSadmix analyses, however, we also looked at patterns of clustering from our data since the method often underestimates K (Gilbert et al., 2012; Janes et al., 2017; Waples & Gaggiotti, 2006). We used ANGSD to compute genetic distances from genotype probabilities and ngsDist (Vieira et al., 2016) for a multiple dimensional scaling analysis (MDS) with the same parameters used for PCA. Results were visualized in the tidyverse (Wickham et al., 2019) with RStudio (Posit team, 2025) and R (R Core Team, 2021).

### 2.6 Mitochondrial genomic phylogeny

We extracted mitochondrial genomic data from WGS reads with the programs BWA MEM v.0.7.17, seqtk, and SAMTools v1.7-1 with the published mitochondrial genome (NC_020311) and mapped them within the program Geneious Prime®2024.0.7. We also aligned genomes with the MAFFT algorithm and tested mitochondrial genome alignments for the best phylogenetic model with Modeltest with AIC (Darriba et al., 2020) within Geneious Prime. We used the program MrBayes v3.2.7 (Ronquist et al., 2012) to estimate a phylogeny of mitochondrial genomes including published data from other members of the Alvinocaridae and other closely related deep-sea shrimp. We ran multiple analyses for 10 million generations, sampled every 1000 generations with six chains, and discarded the first 10% of trees as a burnin. We examined multiple runs to ensure we achieved convergence in Tracer v.1.7.2 (Rambaut et al., 2018). Consensus trees were visualized with TreeViewer v.2.2.0 (Bianchini & Sánchez-Baracaldo, 2023). The mitogenome map was visualized with the CG view server (Stothard & Wishart, 2004, http://cgview.ca).

## 3. Results

### 3.1 Distribution, densities, and habitat preference

The two identified species of Rimicaris had different distributions. We found Rimicaris loihi at five sites on the Izu-Bonin-Mariana (IBM) arc across 1900 km and on Loihi Volcano more than 6000 km to the east (Figure 1A). It was the only species of shrimp on Suiyo, Nikko, and Loihi volcanoes, although the sample size at Loihi was small (7 individuals); however, it is not recorded at seven other IBM vent sites. Rimicaris loihi was the dominant species on NW Rota where, of the 332 specimens confirmed via COI identification, only three were R. cambonae (0.9%). In contrast, R. cambonae was the dominant species in 2023 NW Eifuku collections at 90.3%. However, in the small 2004 sample from the same site, only two of the 25 specimens collected were R. cambonae. Finally, 12 R. cambonae were identified from the West Mata vent field in the Southwest Pacific as the only species present.

R. loihi appeared most abundant on sulphur crusts at all collection sites but were also observed on surfaces of andesitic lavas at NW Rota and among mussels at both NW Eifuku and Suiyo. At NW Eifuku in 2023, R. cambonae co-occurred with R. loihi within mussel beds and on bare rocks.

Repeated exploration of NW Rota vent field between 2004 and 2010 revealed that, as eruption intensity and venting varied over the years, so did the abundance of shrimps (Table 2, Figure 2B–E). For 2004, we estimated average shrimp densities around vents to be 386 m^-2^ (n=6; SD=165); maximum adult densities exceeded 900 m^-2^ in 2009 when shrimps were most abundant (Figure 2B). In the rugged terrain, it was not possible to map the distribution of shrimp species, however we observed that along the known sites, the abundance of shrimps mirrored the venting extent, peaking in 2009 at 45%. The lowest abundances of shrimps were in 2006 during intense eruptions when shrimps were tightly clustered around a few vent openings, and in 2014, when volcanic activity had ceased and hydrothermal venting contracted to the central eruption pit (Table 2; Figure 2E).

**Figure 2.**
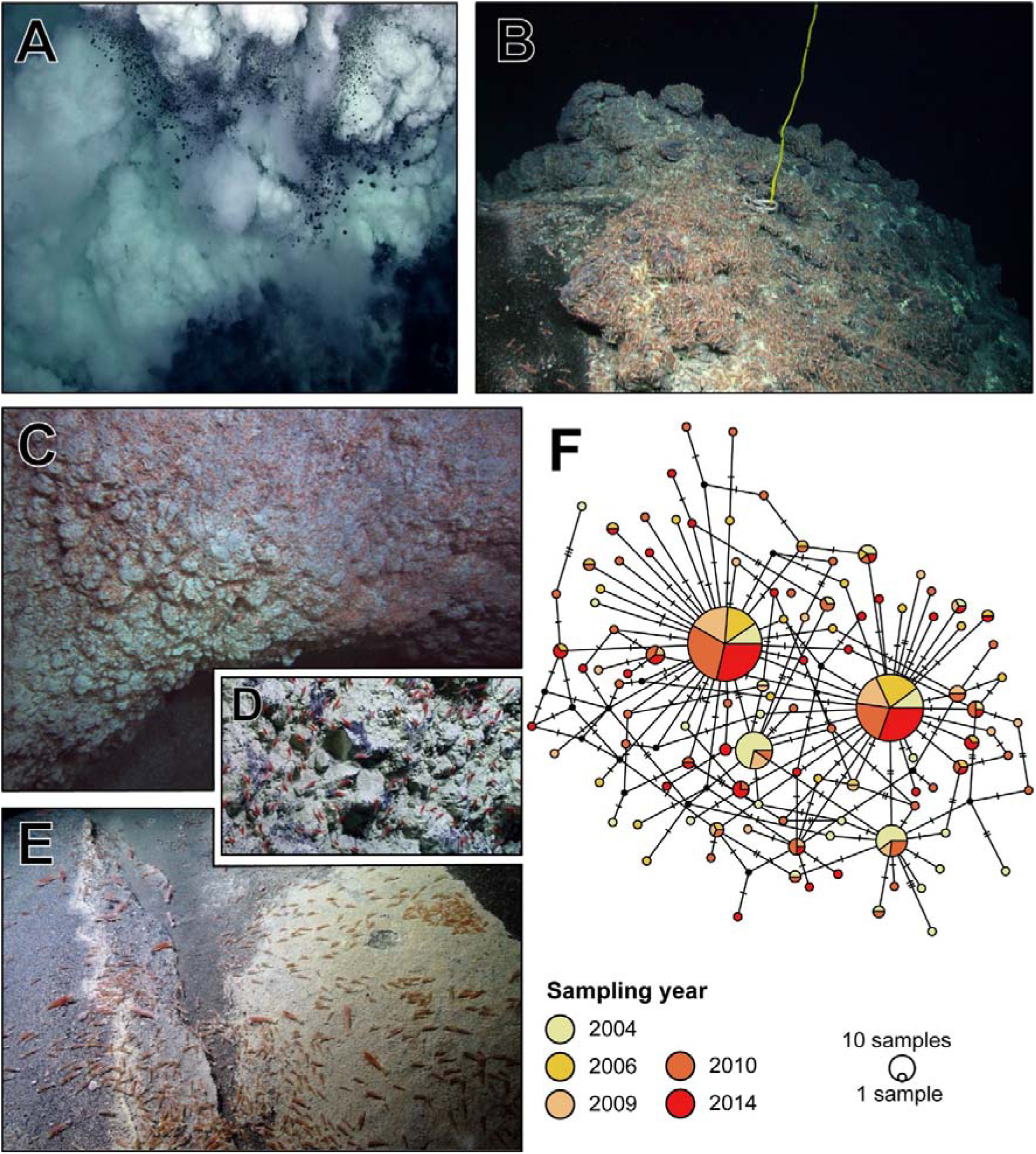
Temporal metapopulation dynamics of Rimicaris loihi at NW Rota between 2004 and 2014. **A.** Eruptive pit with explosive ejection of tephra and sulphur at Brimstone Pit in April 2006. **B.** Abundant shrimps near Marker 109 in April 2009. **C.** New shrimp recruits on sulphur wall in March 2010; red laser dots in the center of image are 10 cm apart. **D.** Close up view of small shrimp recruits on sulphur wall. E. Shrimp in lower abundance in December 2014. **F.** COI haplotype network of Rimicaris loihi from NW Rota (n= 332) detailed per sampling year.

### 3.2 COI haplotype network and genetic diversity

Our Assemble-Species-by-Automatic-Partition (ASAP) analysis to assess putative species revealed two partitions that had equal support (lowest ASAP score of 2.50; Figure S1): individuals fell into either two or three genetic groups. Our morphological identifications of 144 shrimps from NW Eifuku corresponded to two groups; assigned to Rimicaris loihi and R. cambonae. The third COI lineage identified in the three partitions result included distinct R. cambonae haplotypes from NW Rota. Distribution of pairwise differences was bimodal with two distinct Gaussian distributions and showed no overlap (Figure S1). F_ST_ values also supported genetic differentiation for two groups, and possibly a third, with high values between R. cambonae and R. loihi and, to a smaller extent, between R. cambonae from NW Eifuku and from NW Rota (Table S2). The partitions were evident with a relatively low genetic distance (between R. cambonae from NW Eifuku and NW Rota (D_a_= 0.009) compared to the higher genetic distance between R. cambonae and R. loihi (D_a_ = 0.023).

Species delineation was also clear based on the COI haplotype network; it comprised 158 haplotypes from 597 barcoded individuals, of which 134 were assigned to R. loihi and 24 to R. cambonae (Figure 1D). Rimicaris loihi and R. cambonae showed distinct genetic characteristics with a much lower haplotype diversity for R. cambonae compared to R. loihi (H_d_= 0.499 ± 0.052 and H_d_= 0.877 ± 0.011, respectively) and almost three-fold lower nucleotide diversity for R. cambonae (π = 0.00190 and π = 0.00437; Table S3). Similarly, the average number of nucleotide differences was lower for R. cambonae (k= 1.048) than for R. loihi (k= 2.208), a trend that did not appear to be related to sample size differences, with a similarly high value for R. loihi (k = 2.534) when randomly subsampled to the same sample size as R. cambonae. The West Mata population of R. cambonae had higher levels of haplotype diversity (H_d_), nucleotide diversity (π), and average nucleotide differences (k) compared to the northern R. cambonae populations from the IBM arc (Table S3); the southern population also had two intermediate haplotypes, although they are closer to R. cambonae than R. loihi (Figure 1D). Haplotype variability among populations of R. loihi was notably higher than the sympatric R. cambonae, where Suiyo was the most diverse and Nikko was the least diverse (Table S3).

To explore further the possibility of temporal metapopulation dynamics of R. loihi, we estimated a separate haplotype network and summary statistics for NW Rota data only (Figure 2F). Yearly data were similar except for 2006 and 2014 when there was slightly lower nucleotide diversity (π), as well as lower average nucleotide differences (k) (Table S4).

### 3.3 Mitochondrial genome phylogeny

We aligned 26 mitochondrial genomes from R. cambonae and R. loihi recovered from WGS data. Mitochondrial genomic alignments showed the same pattern of divergence between the two species as the COI fragment and differed by 1.5% for the GTR substitution model. The mitochondrial genome of each species had the same gene composition and arrangement as all other published mitochondrial genomes of the Alvinocarididae (Figure 3).

**Figure 3.**
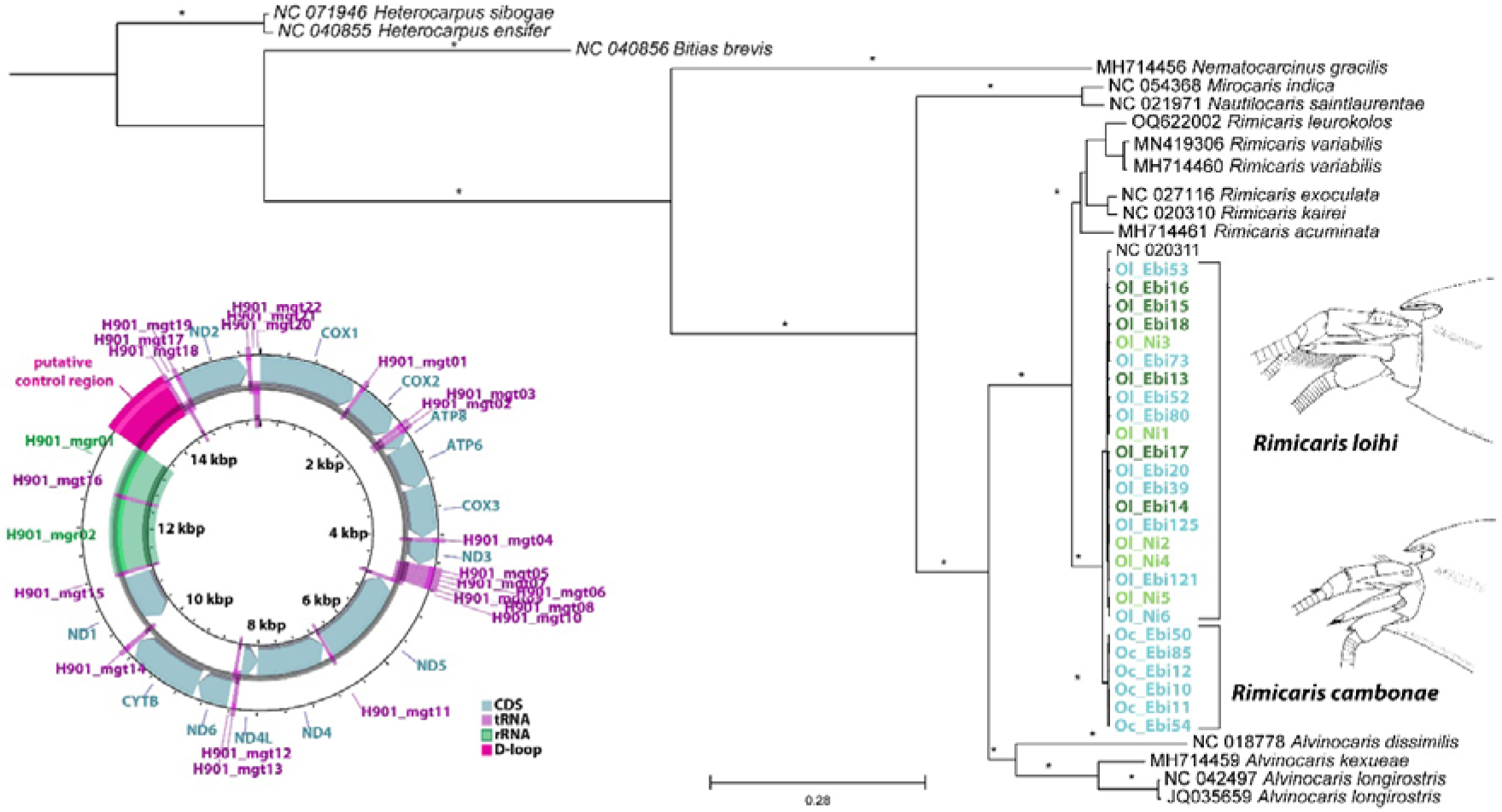
Bayesian estimation of phylogeny of mitochondrial genomic data for Alvinocaridae including R. loihi and R. cambonae samples from Suiyo, Nikko, and NW Eifuku seamounts with the GTR+I+G model for 17,813 sites and a representation of the gene position, type, and order in the mitogenome of R. loihi and R. cambonae. * indicates 1.0 posterior probability.

### 3.4 Whole Genome Shotgun sequencing

Our Whole Genome Shotgun (WGS) sequencing of Rimicaris individuals resulted in 109 million reads that included 19 R. loihi from NW Eifuku, Suiyo, and Nikko seamounts and seven R. cambonae from NW Eifuku (Figure 4A) after we excluded low-coverage individuals. The assembled reference genome from our most successfully sequenced individual of R. cambonae from NW Eifuku (Ebi50) recovered a relatively poor assembly of ∼62.8 m bases with an average coverage of 6x. After we removed scaffolds <1500 bp long the remaining assembly included 1332 scaffolds of 6 million base pairs with an average coverage of 13x and a N50 of 2790 bp.

**Figure 4.**
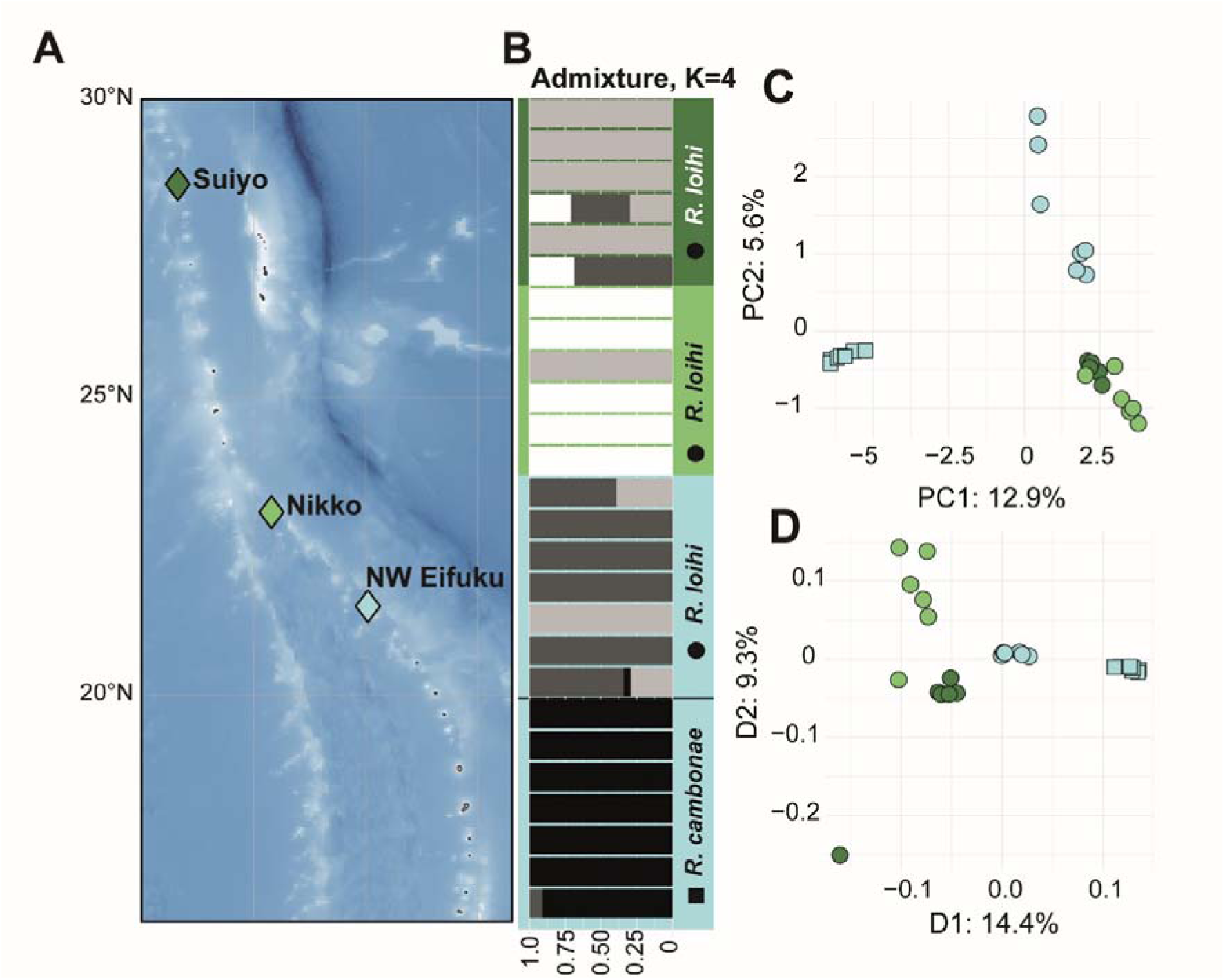
**A.** Map of sampling localities included in population genomic analyses for R. cambonae and R. loihi including Suiyo Seamount (dark green), Nikko Seamount (light green) and NW Eifuku Seamount (blue) including dominant current vectors from Mitarai et al 2016 (arrows). **B.** Admixture plot for four populations (K=4) including 26 individuals for 111002 loci. **C.** Plot of first two axes of principal component analyses (PCA) for 111002 loci. **D.** Plot of first two dimensions for multiple dimensional scaling (MDS) estimates from 126652 loci based on genetic distances for R. cambonae (squares) and R. loihi (circles).

#### 3.4.1 Population Statistics

We estimated summary statistics within each population to examine whether they were in equilibrium (Figure S2). Estimates of summary statistics for R. loihi included 2,175,571 sites from Suiyo, 2,539,205 sites from Nikko, 2,641,049 sites from Eifuku, and 2,837,793 sites from R. cambonae from Eifuku seamounts. The mean Tajima’s D statistic for all populations was similar and slightly positive, although the values were higher for both species from NW Eifuku (T_cam_ = 1.0, T_loh_ = 1.1) than R. loihi from Nikko (T_lohN_ = 0.6) or Suiyo (T_lohS_ = 0.6) they were still below the threshold indicative of balancing selection or strong population contraction. The mean per-site nucleotide diversity ranged from 0.01–0.02 and was lowest in the R. cambonae from NW Eifuku population and highest for the R. loihi population at Nikko seamount. Mean per-site estimates of Waterson’s θ also were relatively low, and ranged from 0.01–0.02 and again was lowest for R. cambonae from NW Eifuku and highest for R. loihi from Nikko seamounts.

#### 3.4.2 Species delineation

We first used genomic data to assess whether nuclear loci were concordant with mitochondrial data and whether they supported R. loihi and R. cambonae as distinct species. After quality filtering, our covariance matrix for PCA included 111,002 sites and revealed population segregation where the majority of differentiation was between R. loihi and R. cambonae (Figure 4). The MDS plot which was based on genetic distances included 126,652 sites and showed that, among R. loihi populations, the one from NW Eifuku was the most closely related to R. cambonae from the same site.

Genotypic admixture plots were estimated with 111,002 sites for two to five populations also revealed differentiation between R. cambonae and R. loihi (Figure S3). The Evanno method showed the most probable number of admixture clusters was four, however the ΔK for two populations was undefined due to a low likelihood variance between K=2 and K=3.

F_ST_ estimates between the species were relatively low, however we detected some subdivision. Global weighted F_ST_ estimates calculated within ANGSD between the populations ranged from 0.103–0.175, where they were lowest between R. cambonae and Suiyo seamount R. loihi and highest between R. cambonae and Nikko Seamount R. loihi (Table S5). F_ST_ estimates calculated with StAMPP were similar to those from ANGSD/realSFS, where they were again low, but all population comparisons were significant (Table S6).

When we compared the demographic histories and possible migration between sympatric populations of R. cambonae and R. loihi from NW Eifuku with the dadi-pipeline we included 141,021 segregating sites with projection sizes of 14 alleles for each species (Figure 5, Table S7). We did not rescale our estimates because we did not have an appropriate mutation rate. We found the most probable model that fit our data was no migration between the species (no_mig, AIC=153626, Figure S4). The next most probable models were divergence with continuous asymmetrical migration (asym_mig, AIC=155126), divergence with different migration rates, with instantaneous population size changes (asym_mig_size, AIC=157445), and divergence with no gene flow, followed by instantaneous size changes with continuous asymmetrical migration (sec_contact_asym_mig_size, AIC=157458, table S7). When we compared allopatric populations of R. cambonae from NW Eifuku and R. loihi from Nikko seamount we included projections sizes of 14 and 12 alleles for each population respectively for 138,269 segregating sites. The most probable model between the allopatric populations was for a split with no migration, then instantaneous size change with no migration (no_mig_size, AIC=144207, Table S8). The next most probable models were divergence with no gene flow, followed by instantaneous size changes with continuous asymmetrical migration (sec_contact_asym_mig_size, AIC=145792), no migration (no_mig, AIC=147041), and divergence with continuous asymmetric migration (anc_asym_mig, AIC=147130, Table S8).

**Figure 5.**
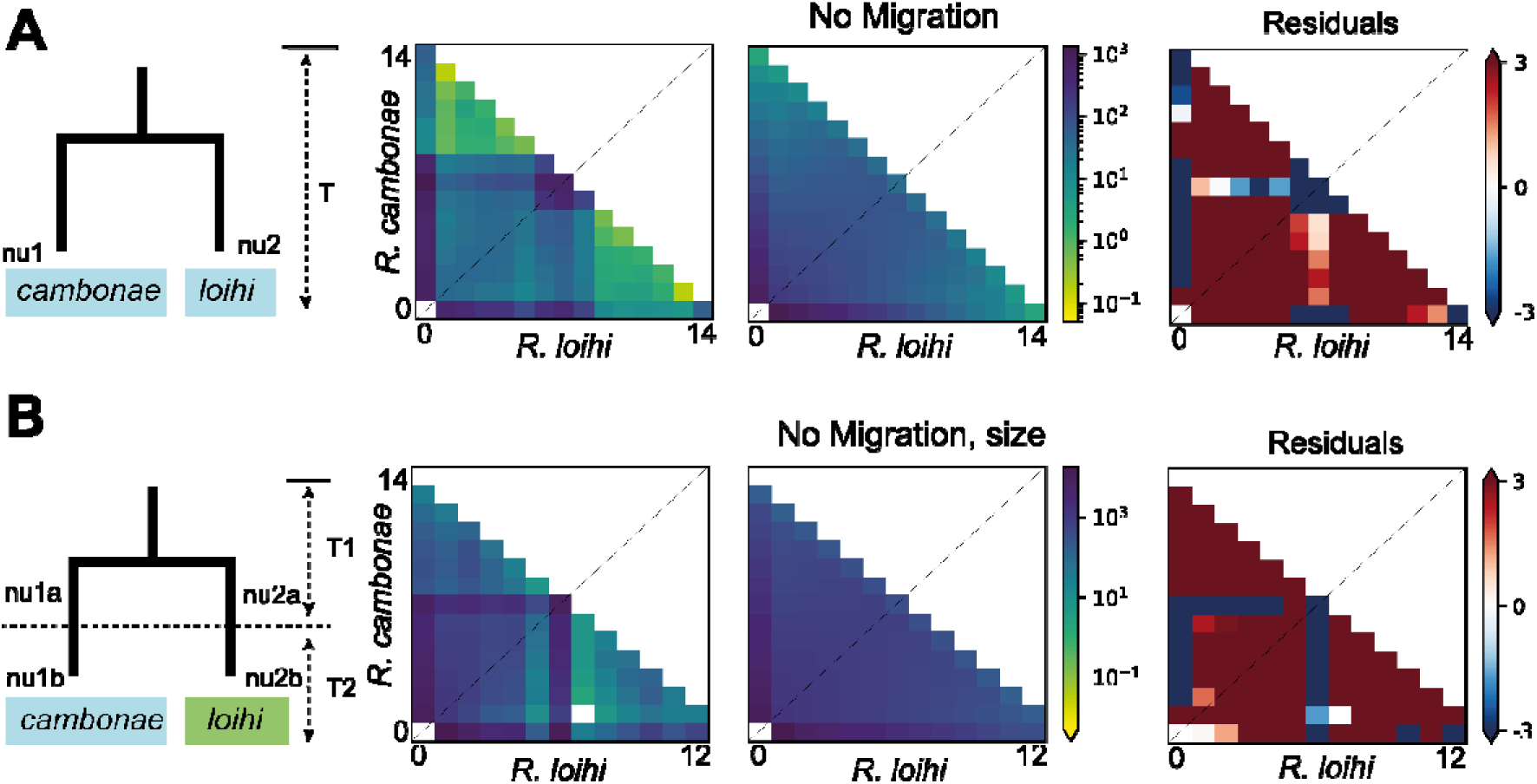
Graphics for the most likely models for dadi_pipeline results and the joint folded allele frequency spectra (AFS), fitted AFS and residuals for **A.** Sympatry: No migration between sympatric populations of seven R. cambonae and seven R. loihi from Eifuku Seamount and **B.** Allopatry: No migration with population size change between allopatric populations of seven R. cambonae and six R. loihi from Nikko Seamount. Cell darkness corresponds to the number of minor alleles in the given bin for both R. loihi and R. cambonae.

#### 3.4.3 Gene Flow within R. loihi

We also found some subdivision among R. loihi populations with MDS and PCA analyses (Figure 4). The NW Eifuku population was the most distinct from Nikko or Suiyo populations, a result reflected by higher F_ST_ values (0.013–0.075, Table S5 and S6). Despite the relative geographic proximity of Nikko and NW Eifuku, there is low probability of larval exchange between them due to ocean current direction (Figure 1, Mitarai et al., 2016). We found low amounts differentiation between Nikko and Suiyo seamounts. The most probable admixture plots for three or more populations also showed that R. loihi from NW Eifuku appeared somewhat divergent and the Suiyo population was a mixture of Nikko and NW Eifuku genotypes (Figures 4 and S3).

Pairwise estimates of demographic histories and migration among three populations of R. loihi from Suiyo, Nikko and NW Eifuku seamounts from dadi_pipeline analyses also remained unscaled because we lacked an appropriate mutation rate. Pairwise analyses based on a folded spectrum between R. loihi populations confirmed patterns of subdivision. We included 136,936 segregating sites with projections for 12 alleles from both Suiyo and Nikko seamounts. We found the most probable model for Suiyo-Nikko interaction was a split with continuous symmetrical migration followed by instantaneous size change (anc_sym_mig_size, AIC=130546, Figure 6, Table S9). The three next most probable models were only slightly less so but involved patterns of admixture instead of migration (Table S9). When we compared 157,151 segregating sites for 12 and 14 allele projections for Nikko and NW Eifuku, we found the most probable model was an island model: vicariance, middle unidirectional discrete admixture event between two drift events (vic_two_epoch_admix, AIC=181429). The next most probable models either involved no migration or a split with continuous symmetric migration, followed by isolation. Lastly, when we compared 128,388 segregating sites for 12 and 14 allele projections the most probable model for comparisons between Suiyo and NW Eifuku also had no migration but was a founder event with a middle unidirectional discrete admixture event (between two drift events) (founder_nomig_admix_two_epoch, AIC=129489). However, two of the three next most probable models both had asymmetric migration from Suiyo into NW Eifuku populations (Table S9).

**Figure 6.**
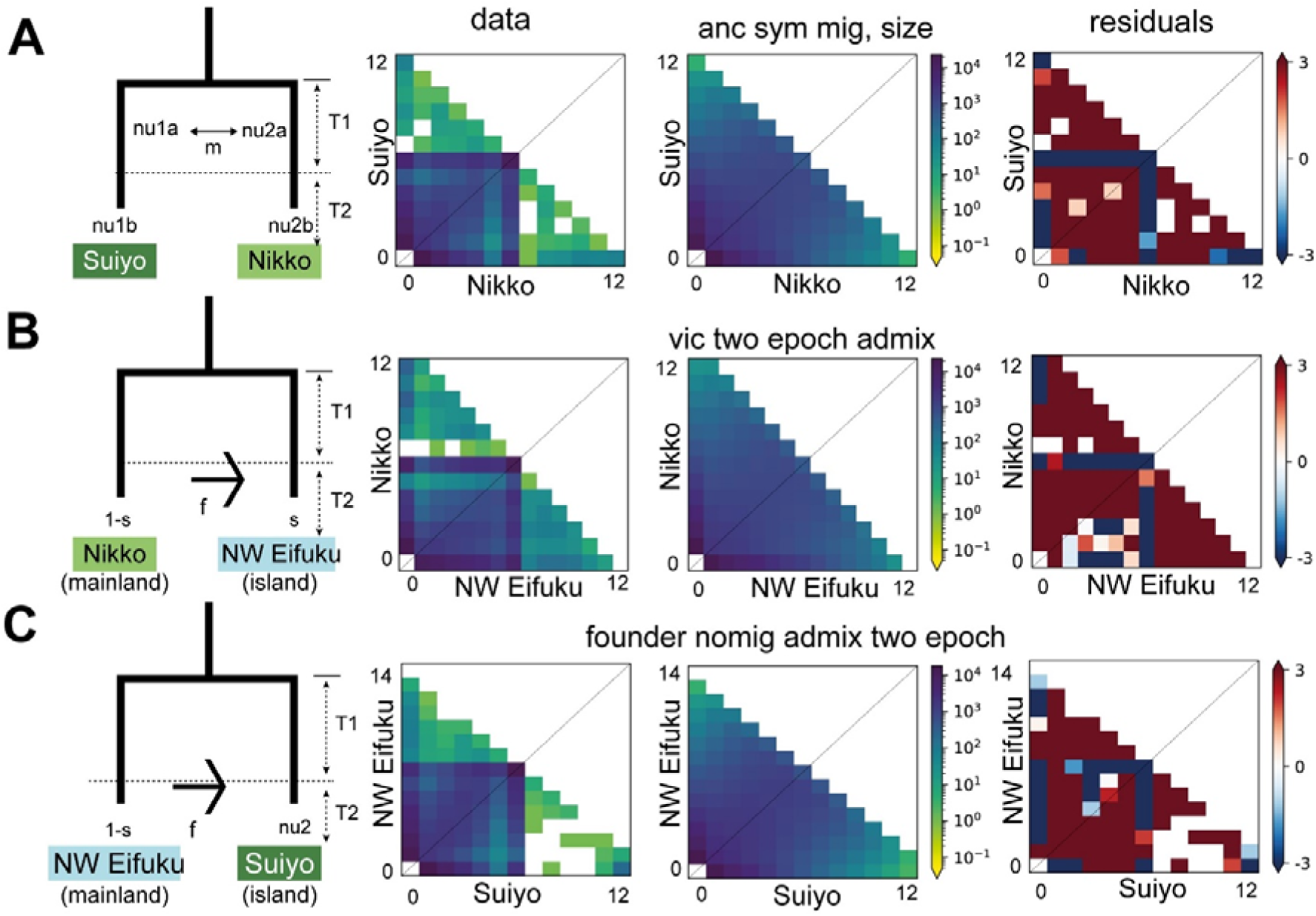
Graphics of the most likely models from dadi_pipeline results and the joint folded allele frequency spectra (AFS), fitted AFS, and residuals within R. loihi for **A.** Ancestral symmetrical migration with size changes between Suiyo and Nikko Seamount, **B.** Island model of vicariance, middle unidirectional discrete admixture event between two drift events between Nikko and NW Eifuku Seamount and **C.** The island model for a founder event with a middle unidirectional discrete admixture event between two drift events for Eifuku and Suiyo seamount. Cell darkness corresponds to the number of minor alleles in the given bin for both populations.

**Figure 7.**
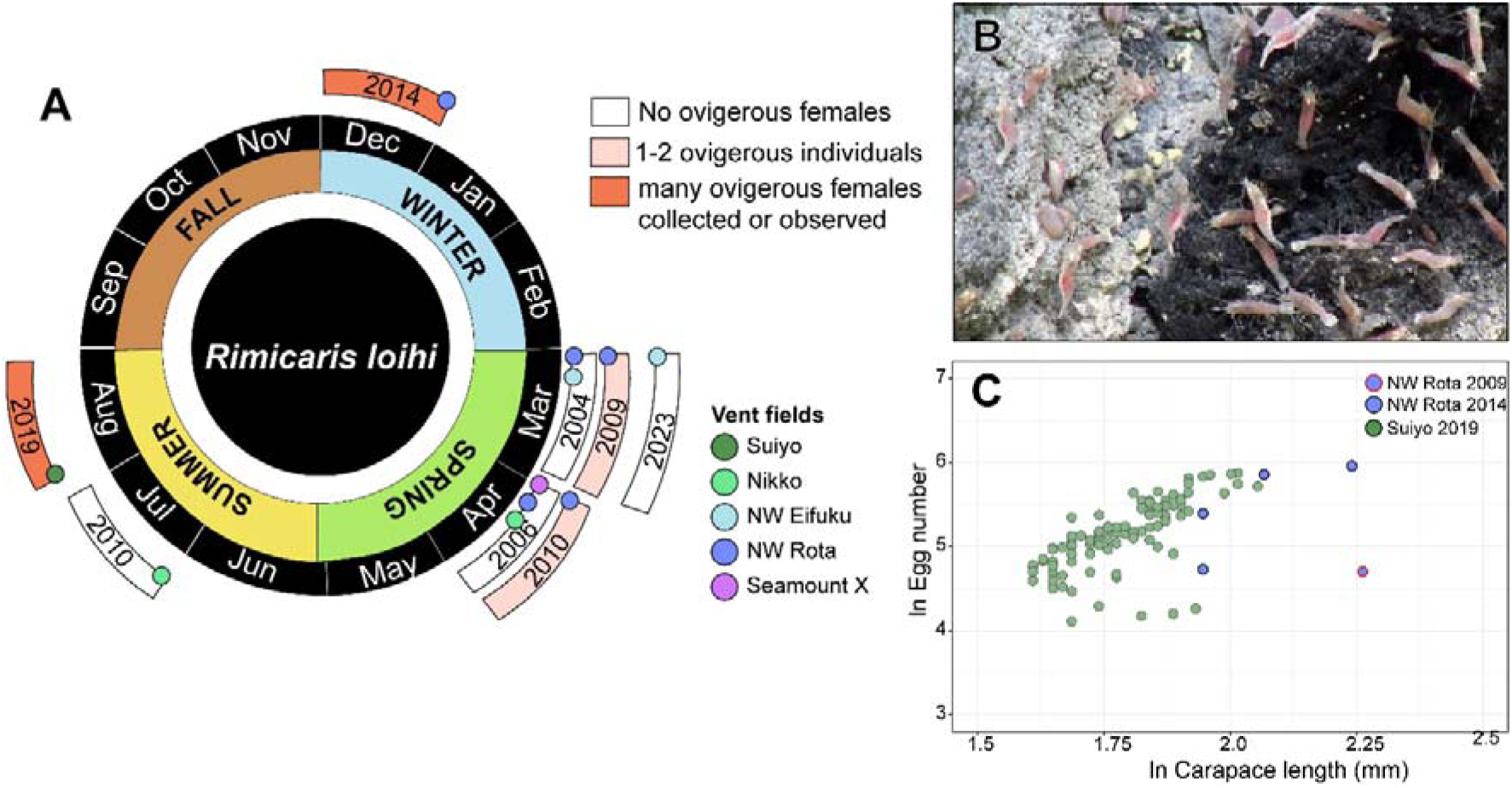
Reproductive characteristics of Rimicaris loihi. **A.** Diagrams summarizing the occurrence of R. loihi ovigerous females over sampling realized between 2004 and 2019. **B.** Numerous ovigerous females (reddish colour below the abdomen) observed, but not collected, at NW Rota in December 2014. **C.** Relative fecundity of R. loihi displayed by log_e_-transformed number of eggs with log_e_-transformed carapace length (CL). Data from Suiyo are from Methou et al. (2023). Data from NW Rota are from this study.

### 3.4 Population sex ratio and reproductive traits

The sex ratio of R. loihi from NW Rota was significantly female skewed for each year (χ^2^ = 25.64 – 259.69, p < 0.001; Table 1), as previously observed in the Suiyo population (Methou et al., 2023). A similar trend occurs for the sex ratio of R. loihi from NW Eifuku, but was not statistically supported (χ^2^ = 1.33, p > 0.05) probably due to a low sample size (Table 1). Conversely, the sex ratio of R. cambonae from NW Eifuku did not significantly deviate from 1:1 (χ^2^ = 1.09, p > 0.05; Table 1). Despite large sampling sizes, ovigerous females of R. loihi were nearly absent from populations collected at NW Rota in March (2004 and 2009) and April (2006 and 2010) with only two ovigerous females out of 408 females in these years (Figure 4A). Similarly, no ovigerous females were collected among the 72 females of R. cambonae from NW Eifuku in March 2023 (Figure 6A). However, four ovigerous females out of 380 females were collected at NW Rota in December 2014 (Figure 6A) and larger numbers were observed peripheral to venting (but not sampled; Figure 6B). With 110 to 387 eggs per brood (11.4 to 44.4 eggs mm^−1^), relative fecundities of R. loihi ovigerous females from NW Rota were in the range reported at Suiyo with 61 to 356 eggs per brood (10.2 to 49.2 eggs mm^−1^), (Figure 6C) (Methou et al., 2023).

**Table 1.** Population characteristics with sex ratio of Rimicaris cambonae and Rimicaris loihi from NW Rota, Suiyo and NW Eifuku. ^(1)^ Data from Suiyo are from Methou et al. (2023). M: males, F: females; Significance levels: NS: > 0.05, *: < 0.05, **: < 0.01, ***: < 0.00.

| Vent field | Year | Species | M | F | M:F | $\chi^2$ | *p* |
| --- | --- | --- | --- | --- | --- | --- | --- |
| NW Rota | 2009 | *Rimicaris loihi* | 119 | 211 | 0.56 | 25.65 | *** |
| NW Rota | 2010 | *Rimicaris loihi* | 395 | 797 | 0.49 | 135.57 | *** |
| NW Rota | 2014 | *Rimicaris loihi* | 47 | 380 | 0.12 | 259.69 | *** |
| Suiyo<sup>(1)</sup> | 2019 | *Rimicaris loihi* | 145 | 235 | 0.62 | 22.94 | *** |
| NW Eifuku | 2023 | *Rimicaris loihi* | 4 | 8 | 0.5 | 1.33 | NS |
| NW Eifuku | 2023 | *Rimicaris cambonae* | 60 | 72 | 0.83 | 1.09 | NS |

**Table 2.** History of observed volcanism and venting on NW Rota. The relative extent of the shrimp populations is assessed by estimating the % of ROV tracks in which shrimp were greater than 20 individuals/m^2^. ^(1)^Embley et al. 2006; ^(2)^Chadwick et al. 2008; ^(3)^Chadwick et al. 2012^; (4)^Schnur et al. 2014.

| Year | Volcanic state | Hydrothermal State | Shrimp population |
| --- | --- | --- | --- |
| 2004 | Volcanic vent in a deep pit with surging sulphurous plume and ejecta; mass wasting on volcano flank.<sup>(1)</sup> | 13 discrete low temperature vents and seeping around rim of volcanic pit. | Extensive: dense at vents and scattered across summit; 37% |
| 2006 | Explosive eruptions from pit observed after sidewall collapse; ejecta volume to 100 m<sup>3</sup>/hr.<sup>(2)</sup> | Discrete sites fewer; seeping through sediments near pit; eastern scarp more focussed. | Restricted: scattered near pit; concentrated at three sites; 12%. |
| 2009 | Up to 47 m of tephra accumulated since 2004<sup>(3)</sup>; ejecta evident. | Many discrete venting sites <50°C broadly distributed across summit; venting through sediments. | Very extensive: dense at vents and scattered across summit; 45% |
| 2010 | Recent eruption reshaped the summit<sup>(3)</sup>; now several volcanic orifices | Loss of several vents in landslide; exposed walls covered with sulphur; | Very extensive: dense at vents; scattered on summit and on new exposed walls; 40%. |
| 2014 | No eruptive activity<sup>(4)</sup>; likely quiescence >1 year. | No or negligible activity at all former sites; only notable temperature anomaly around pit. | Restricted mostly to one site; other locations limited or abandoned; 6%. |

## 4. Discussion

Nuclear genomic data confirmed previous results based on the mitochondrial COI fragment and morphology suggesting that Rimicaris loihi and Rimicaris cambonae constitute two separate species (Methou, Chen, et al., 2024). This segregation into two distinct species was further supported by our demographic modelling analyses that indicated strict isolation with no migration as the most probable model; thus, gene flow has likely stopped between R. cambonae and R. loihi despite their sympatric distribution.

Genetic structure is also present among the different populations of R. loihi along the Izu-Bonin-Mariana (IBM) arc. Yet, demographic histories supported ancient migration or secondary contact as the most probable model, suggesting that some level of gene flow is maintained between populations from Suiyo to NW Eifuku. Patterns of genetic structure for R. loihi populations was not simply a result of distance since we observed a greater genetic similarity between the more geographically distant populations of Suiyo and Nikko than with NW Eifuku, which is closer to Nikko. This pattern contrasts with most vent endemic species in the West Pacific that exhibit high genetic homogeneity within-basin or within-arc structures (Breusing et al., 2021; Poitrimol et al., 2022; Tran Lu Y et al., 2022, 2025). Only the copepod Stygiopontius lauensis, whose geographic distribution is constrained within the Lau Basin, shows similar patterns of structured populations at a regional scale (Diaz-Recio Lorenzo et al., 2024). However, such structuration at a small spatial scale on the IBM arc was unexpected for R. loihi, especially given its broad distribution outside the region, including to Loihi Seamount near Hawaii. This broad geographic distribution also applies to its sister species, since previous reports of R. loihi at West Mata volcano (Resing et al., 2011) actually represent R. cambonae according to our barcoding analyses.

Alvinocaridid shrimps are generally considered as good dispersers with high genetic connectivity over long distances as observed in many species (Methou, Ogawa, et al., 2024; Thaler et al., 2014; Yahagi et al., 2015; Zhou et al., 2022). For example, R. exoculata lacks genetic structure over 7100 km, likely supported by Mid-Atlantic Ridge ocean circulation promoting connectivity (Teixeira et al., 2012). Morphological features of their hatched larvae including R. loihi also supports a long pelagic larval duration (PLD) (Hernández-Ávila et al., 2015; Methou et al., 2023). However, calculated probabilities for larval exchange among our sites are low (Mitarai et al. 2016). The dominant upper ocean currents in this region are across, not along, the Arc in an east to west direction (Kendall & Poti, 2014), thus most larvae are advected away from the Arc although some may be retained.

The genetic structuration among regional R. loihi populations is likely also driven the particular geological context of the IBM arc and enforced by ocean currents. Several hydrothermal vent fields on the IBM volcanic arc are characterized by high geological instability (Embley et al., 2007; Lupton et al., 2006). At NW Rota, our observations over 10 years revealed repeated eruptions with significant variation in the extent of venting activities. Shrimp abundance over the years reflected these changes of venting that also affected their genetic diversity as the NW Rota population declined or expanded. This strong instability of habitat availability may lead to repeated local contraction/extinction of R. loihi at some sites on a temporary basis; rupture of connecting nodes could isolate shrimp populations to less connected areas, such as the northern or southern parts of the IBM arc (Mitarai et al., 2016). This limited connectivity at a regional scale may explain the absence of R. loihi at several IBM vent sites (Brunner et al., 2022) and the apparent lack of R. cambonae (or cf. loihi) beyond the northern Tonga-Tofua volcanos of the SW Pacific. These shrimps also appear to have specific habitat requirements linked to the volcanic emissions or sulphur substratum that characterize all the sites colonized by both species.

Sea level changes during glaciation periods may also have contributed to the temporary isolation of R. loihi populations. During the Last Glacial Maximum, sea-level decrease could have transformed shallowest arc volcanoes into subaerial volcanoes, eliminating chemosynthetic habitats that could serve as potential stepping stones for vent endemic species like Rimicaris shrimps. This contrasts with what we know from terrestrial archipelagos where sea level drop may have reduced isolation between some islands (Fernández-Palacios, 2016; Weigelt et al., 2016). Periods of relative quiescence in volcanism and venting associated with episodic delivery of magma in the rising melt (Stern et al., 2003) may also have prevented the establishment of chemosynthetic fauna during these intervals.

The same mechanisms that drive geological instability at a local scale or related to ocean currents and sea-level changes may also have promoted speciation between R. cambonae and R. loihi by isolating two ancestral populations for a sufficient period of time. The genetic similarity between R. cambonae and R. loihi is closest on NW Eifuku; initial separation may have occurred at this site or between ancestral population not in our analyses. This could include populations from the southern part of the Mariana arc (i.e., NW Rota) or from the more remote populations of West Mata in the Tonga arc or Loihi seamount near Hawaii. Given the predominant eastward North Pacific Drift, the latter site is mostly likely a sink, not a source, location. Larval dispersal among volcanic arc systems or back-arc basins in the West Pacific is inferred to be infrequent and directional, following oceanic currents (Mitarai et al., 2016; Tunnicliffe et al., 2023). In particular, transit is deemed highly improbable between south and north hemispheres as larvae transport is inhibited by the strong Equatorial Counter Current and Equatorial Undercurrent that both flow eastward in the upper 300 m with little water movement below (Constantin & Johnson, 2015). Thus, relatively few hydrothermal vent species distributions cross the equator (Tunnicliffe et al., 2023). A rare circumstance of long-distance dispersal from either the IBM southward or the Tonga-Tofua Arc northward could have established an outpost population in the distal arc; we cannot determine the direction. Subsequent geological instability in both volcanically active regions may have affected shrimp population sizes and genetic diversities to different extents in both source and recipient sites. The current sympatric distribution of both R. loihi and R. cambonae in the Mariana Arc likely results from secondary contact by another rare larval dispersal event from south to north. This scenario of speciation by allopatry postulates that genetic inflow via larval dispersal between the two regions was disrupted for a sufficiently long period to generate an isolation phase that would persist and be maintained in sympatry.

Genetic isolation between sympatric sister species could be maintained by strong reproductive barriers. Differences in reproductive mode (egg laying versus brooding) strengthens reproductive isolation between the syntopic sister species of intertidal snails Littorina saxatilis and L. arcana (Stankowski et al., 2020). In coastal isopods of the Jaera species complex, sexual isolation between co-occurring species is reinforced by preferential mate choice even with some level of hybridization and no influx from parental populations (Ribardière et al., 2017, 2021). Allochronic isolation with different reproductive seasonality can also maintain genetic differentiation as reported in Mediterranean pine moths (Santos et al., 2007). However, our dataset on the reproductive characteristics of R. loihi and R. cambonae remains inconclusive on the possible existence of such barriers to reproduction. As with most carideans (Correa & Thiel, 2003), alvinocaridid shrimps are all gonochoric species that mate and brood their eggs under their abdomen until larval release (Methou, Chen, et al., 2022; Methou et al., 2023). A wide variety of reproductive cycles occur in this family, including aperiodic or periodic brooding patterns, with or without synchronicity to seasonal variations (Copley & Young, 2006; Methou, Chen, et al., 2022; Methou et al., 2023). The presence of numerous ovigerous individuals of R. loihi both in August and December – despite a notable and repeated absence in March/April – does not point to a periodicity distinct from its sister species. We did find that populations of R. loihi from the Mariana Arc, as at Suiyo (Methou et al., 2023), were largely biased toward females whereas the proportion of males and females was roughly equal for R. cambonae. This discrepancy between the sex ratios of the two species could indicate different mating choice, as variation in adult sex ratios can strongly influence breeding systems and mate choice in many animal groups (Grant & Grant, 2019; Székely et al., 2014). Yet assortative mating between the two Rimicaris species remains to be demonstrated. Further studies using larger genomic dataset across the entire range of the two species as well as investigations on their trophic and reproductive ecology will be required to clarify mechanisms leading to their speciation.

## 5. Conclusion

Rimicaris loihi and R. cambonae are two distinct, recently diverged species living in sympatry throughout a part of their geographic range. Hybridisation does not appear to occur between the two species. Analysing genetic data from shrimps collected from a single vent over a decade revealed variability in genetic diversity that correlates with local eruption events, suggesting that population dynamics of these shrimps are impacted by their unstable habitats. Although our demographic analyses remain inconclusive as to how speciation occurred among these two species, we propose allopatric divergence and a secondary contact by a long-distance dispersal event as the most likely scenario. The geographic overlap and recent divergence of this species pair makes them an interesting case study for a future exploration into speciation mechanisms that may provide insights to how species arise in insular habitats overall. Our findings also have an important implication on conservation, showing widespread distribution does not equate to resilience. Local disturbances such as volcanic eruptions or anthropogenic impacts like deep-sea mining might not cause these species to go extinct but can exterminate genetically distinct populations, with limited potential for recovery via recolonization.

## Supporting information

Population_dataset

## Acknowledgments

We thank Dr. Craig Moyer for providing shrimp specimens from Loihi Seamount. We also thank Joe Jones and Lara Puetz for their work on genetic barcoding for some specimens. Captain and crew of R/Vs Yokosuka, Kaimei, and Natsushima are gratefully acknowledged for their continued support of our scientific activities during the cruises YK19-10, KM23-E05, and NT10-13 Leg 2, respectively. We extend this to the pilots and technical teams of HOVs Shinkai 6500 and Pisces V, as well as ROVs Hyper-Dolphin, KM-ROV, Jason-2, and ROPOS. This is a GACHINKO Cruise Episode I (19-10) output, and we thank the GACHI-participants for their help in sorting shrimp specimens on-board. We thank the cruise chief scientists Ken Takai (JAMSTEC; YK19-10, KM23-E05) and Shinji Tsuchida (JAMSTEC; NT10-13 Leg 2), as well as Shinsuke Kawagucci (JAMSTEC) whose successful cruise proposal led to the sampling on the Mariana volcanic arc during cruise KM23-E05. Sampling during cruise KM23-E05 within the Islands Unit of the Mariana Trench Marine National Monument of the United States of America was carried out under the special use permit SUP 12542-23001 (MSR U2022-047). We are grateful for collections during several cruises in the NOAA Ring of Fire series of the Vents Program on the Mariana Volcanic Arc. Chief Scientists Bob Embley, Bill Chadwick, and Dave Butterfield were critical to the output.

## Funding

PM was supported by ISblue project, Interdisciplinary graduate school for the blue planet (ANR-17-EURE-0015) and co-funded by a grant from the French government under the program "Investissements d’Avenir" embedded in France 2030. PM also benefited from French State aid managed by the National Research Agency under France 2030: ANR-22-POCE-0007. VT was supported by NSERC Canada and the Canada Research Chairs programme. Whole Genome Shotgun Sequencing was funded by the David and Lucile Packard Foundation at MBARI.

## Data Accessibility & Benefit-Sharing

### Data Accessibility

All COI barcode sequences have been deposited in GenBank under accession numbers DQ328819 – DQ328837 and PQ643496 – PQ643653 (see Table S10 for details on haplotype numbers and localities with associated individual ID). Raw sequences of the WGS sequencing dataset are available in the NCBI SRA repository with the BioProject ID PRJNA1197504 (Accession numbers: SAMN45807939 – SAMN45807966). Details information on sampling localities, sampling date, sex, life stages and sizes of individuals analysed for population sex ratio and reproduction are compiled in a table sheet named “Population_dataset” in supplementary material.

### Benefit-sharing

Faunal collections were conducted with the necessary authority permissions of the relevant countries. Sampling in the Japanese exclusive economic zone (EEZ) and the Mariana EEZ were respectively conducted by Japanese government and American government research vessels. Permission for sampling in the Tongan EEZ was issued by the Kingdom of Tonga. Research animals were invertebrate caridean shrimps and no live experiments with animals were conducted in this study.

## Notes

### Competing Interest Statement

The authors have declared no competing interest.

### Summary of Updates

Some filtering steps of the WGS analyses were revised which changes some of the results and conclusions from the previous version of the manuscript.

