## Supplementary Material for "A tale of two shrimps – Speciation and demography of two sympatric shrimp species from hydrothermal vents"

### Supplementary Figures

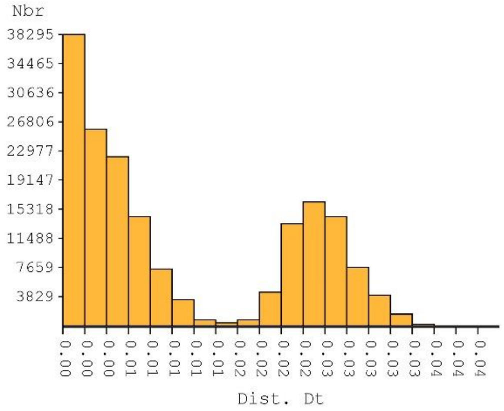


| Nb of subsets | asap-score | P-val (rank) | W (rank) | Treshold dist. |
|---|---|---|---|---|
| 3 | 2.50 | 9.94e-02 (2) | 5.69e-07 (3) | 0.009489 |
| 2 | 2.50 | 2.12e-01 (4) | 1.66e-06 (1) | 0.011931 |
| 5 | 4.00 | 8.02e-02 (1) | 1.43e-07 (8) | 0.006752 |
| 8 | 4.50 | 2.34e-01 (6) | 4.76e-07 (4) | 0.004397 |
| 167 | 5.50 | 1.76e-01 (3) | 8.96e-08 (8) | 0.000682 |
| 32 | 5.50 | 3.37e-01 (6) | 3.34e-07 (5) | 0.002973 |
| 142 | 7.00 | 5.13e-01 (8) | 2.58e-07 (6) | 0.001536 |
| 33 | 9.50 | 3.95e-01 (7) | 6.91e-08 (12) | 0.002345 |
| 143 | 10.50 | 8.20e-01 (19) | 5.74e-07 (2) | 0.001415 |
| 7 | 11.00 | 7.27e-01 (11) | 7.76e-08 (11) | 0.005128 |

**Figure S1.** Species delimitation of *Rimicaris cambonae* and *Rimicaris loihi* using Assemble Species by Automatic Partitioning (ASAP) with the Kimura 80 substitution model. **A.** Histogram of distances. **B.** Two-best partitions and their associated metrics (number of subsets, asap-score, p-values of partitions and associated ranks, W (rank): width of the barcode gap, threshold distance).

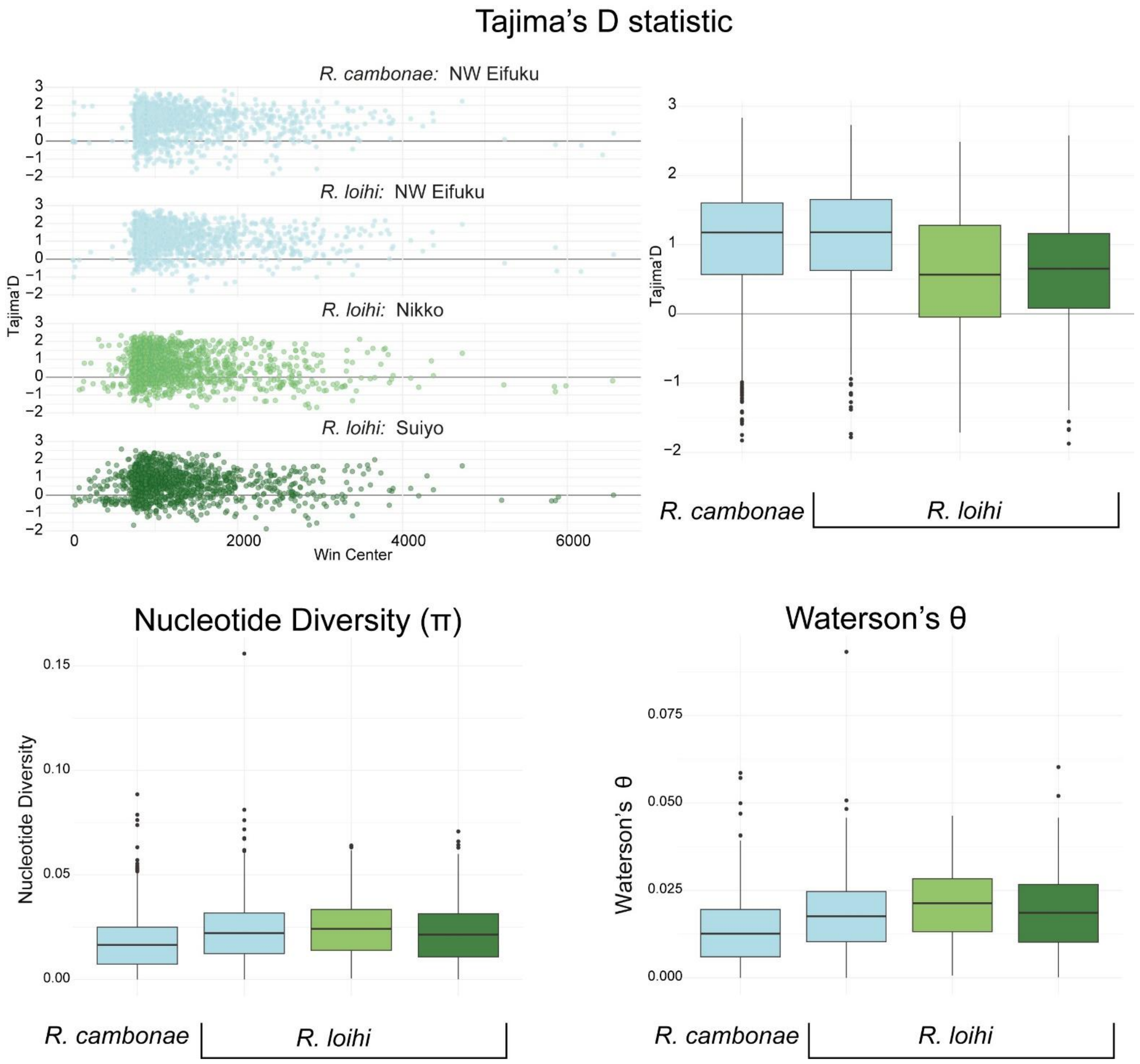


**Figure S2.** Summary statistics of whole genome shotgun sequencing (WGS) with estimates for (**A. B.**) Tajima's D (**C.**) per-site nucleotide diversity and (**D.**) Watterson's theta for *R. cambonae* from NW Eifuku (light blue), and for *R. loihi* from NW Eifuku (light blue),Nikko (light green), and Suiyo Seamounts (dark green).

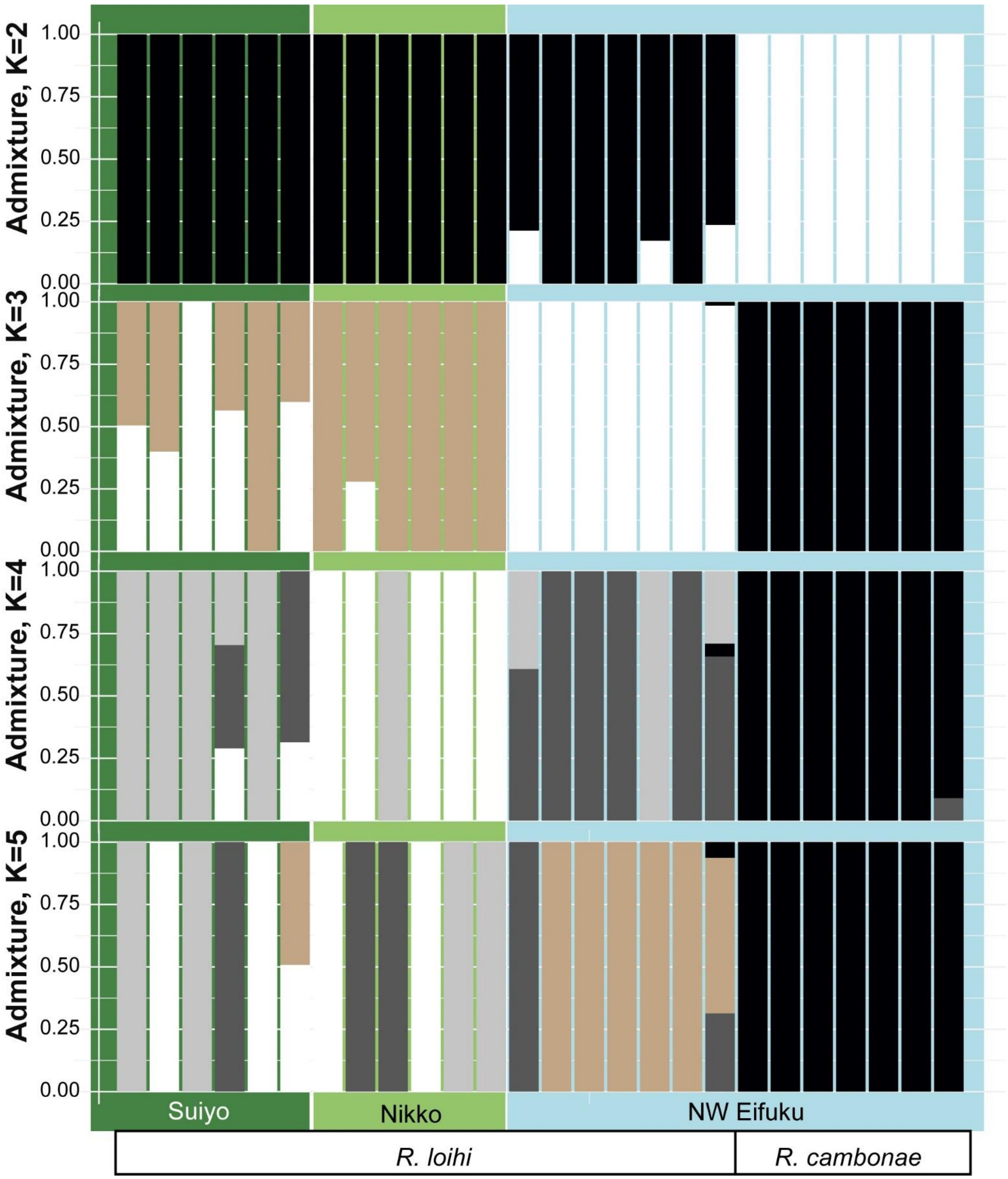


**Figure S3.** Admixture plots for two (K=2) three (K=3), four (K=4) and five (K=5) populations including 26 individuals for 111002 loci.

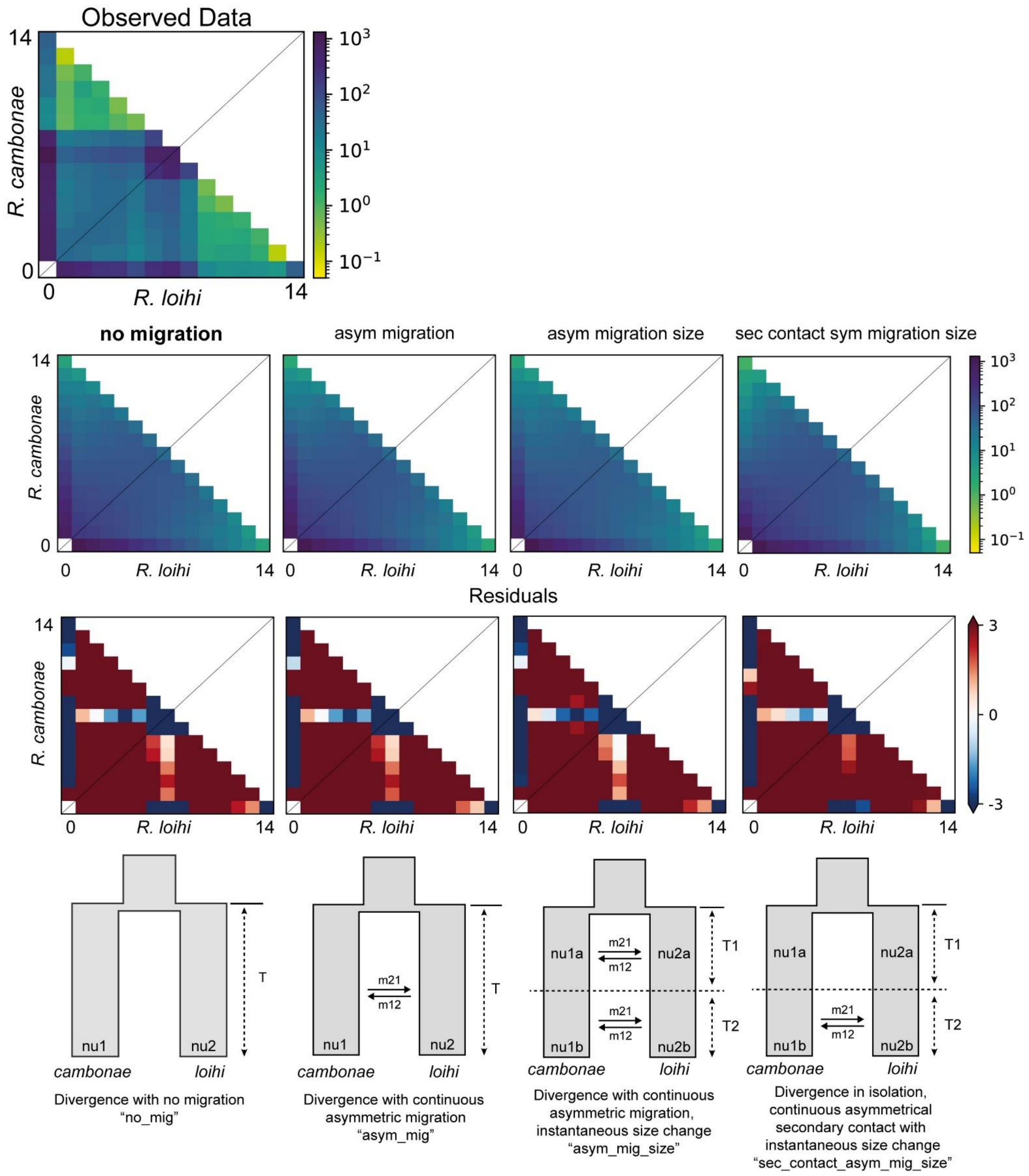


**Figure S4.** Joint folded allele frequency spectra (AFS) for observed (top row) and fitted (middle row) for seven individuals each of *R. loihi* (x-axis) to *R. cambonae* (y-axis) from Eifuku seamount for the four most probable models (bottom row) for dadi analyses. 'No migration' was the most probable model (AIC=153626) and the AFS and residuals are plotted in comparison to the other models. The next most probable models were 'asym_mig' (AIC=155126), 'asym_mig_size' (AIC=157445), and 'sec_contact_asym_mig_size' (AIC=157548). Cell darkness corresponds to the number of minor alleles in the given bin for both *R. loihi* and *R. cambonae*

### Supplementary Tables

| Location | Latitude | Longitude | Depth | Expedition ID | ROV/HOV | Year | Month | COI primers |
|---|---|---|---|---|---|---|---|---|
| **Loihi** | 18.9167 | -155.2667 | 980m | n/a | *Pisces V* | 2004 | | COI-1f/1r[1] |
| **Suiyo** | 28.5716 | 140.6426 | 1376m | YK19-10 | *HOV Shinkai 6500* | 2019 | August | CariCOI-1F/1R[2] |
| **Nikko** | 23.0662 | 142.3166 | 425-460m | MGLN02MV | *ROV Jason-2* | 2006 | April | COI-1f/1r[1] |
| | | | | NT10-13 | *ROV Hyper-Dolphin* | 2010 | July | CariCOI-1F/1R[2] |
| **NW Eifuku** | 21.4875 | 144.0412 | 1560-1640m | TN167 | *ROV ROPOS* | 2004 | March | COI-1f/1r[1] |
| | | | | KM23-05 | *KM-ROV* | 2023 | March | CariCOI-1F/1R[2] |
| **NW Rota** | 14.6009 | 144.7775 | 517-630m | TN167 | *ROV ROPOS* | 2004 | March | COI-1f/1r[1] |
| | | | | MGLN02MV | *ROV Jason-2* | 2006 | April | COI-1f/1r[1] |
| | | | | TN-232 | *ROV Jason-2* | 2009 | April | COI-1f/1r[1] |
| | | | | KM1005 | *ROV Jason-2* | 2010 | March | COI-1f/1r[1] |
| | | | | RR1413 | *ROV Jason-2* | 2014 | December | COI-1f/1r[1] |
| **Seamount X** | 13.2528 | 144.0197 | 1305m | MGLN02MV | *ROV Jason-2* | 2006 | April | COI-1f/1r[1] |
| **West Mata** | -15.0950 | -173.7500 | 1208m | TN-234 | *ROV Jason-2* | 2009 | May | COI-1f/1r[1] |

**Table S1.** Summary of vent fields sampled for this study by expeditions. For each expedition, details of COI primers used for the barcoding dataset are also given. [1] This study. [2] (Methou et al. 2020)

| | Fst | Da |
|---|---|---|
| ***Rimicaris spp.*** | | |
| *R. cambonae* vs *R. loihi* all populations | 0.88578 | 0.02427 |
| *R. cambonae* vs *R. loihi* NW Rota | 0.86495 | 0.02015 |
| *R. cambonae* vs *R. loihi* NW Eifuku | 0.89245 | 0.02319 |
| ***Rimicaris cambonae*** | | |
| NW Eifuku vs NW Rota | 0.91223 | 0.01078 |
| NW Eifuku vs West Mata | 0.13605 | 0.0006 |
| NW Rota vs West Mata | 0.70916 | 0.00964 |
| ***Rimicaris loihi*** | | |
| Loihi vs Suiyo | -0.02797 | -0.00011 |
| Loihi vs Nikko | -0.02864 | -0.00011 |
| Loihi vs NW Eifuku | 0.00373 | 0.00002 |
| Loihi vs NW Rota | 0.04244 | 0.0002 |
| Loihi vs Seamount X | -0.03571 | -0.00017 |
| Suiyo vs Nikko | -0.00277 | -0.00001 |
| Suiyo vs NW Eifuku | 0.02193 | 0.0001 |
| Suiyo vs NW Rota | 0.0635 | 0.00029 |
| Suiyo vs Seamount X | 0.01346 | 0.00007 |
| Nikko vs NW Eifuku | 0.02394 | 0.00011 |
| Nikko vs NW Rota | 0.06658 | 0.00031 |
| Nikko vs Seamount X | -0.01228 | -0.00006 |
| NW Eifuku vs NW Rota | 0.00341 | 0.00002 |
| NW Eifuku vs Seamount X | 0.03597 | 0.00021 |
| NW Rota vs Seamount X | 0.06141 | 0.00035 |

**Table S2.** Pairwise Fst values and raw genetic distance (Da) among populations of *Rimicaris cambonae* and *Rimicaris loihi* based on partial COI haplotype sequences.

| | N | S | h | Hd | π | k |
|---|---|---|---|---|---|---|
| ***Rimicaris cambonae*** | | | | | | |
| NW Eifuku | 125 | 15 | 15 | 0.406 ± 0.056 | 0.00085 | 0.0514 |
| NW Rota | 3 | 1 | 2 | 0.667 ± 0.314 | 0.00120 | 0.667 |
| Seamount X | 1 | - | - | - | - | - |
| West Mata | 12 | 20 | 10 | 0.955 ± 0.057 | 0.00703 | 4.379 |
| All populations | 141 | 29 | 23 | 0.491 ± 0.052 | 0.00191 | 1.056 |
| ***Rimicaris loihi*** | | | | | | |
| Loihi | 7 | 6 | 5 | 0.857 ± 0.137 | 0.00340 | 2.0 |
| Suiyo | 23 | 25 | 20 | 0.984 ± 0.019 | 0.00608 | 4.119 |
| Nikko | 46 | 24 | 19 | 0.805 ± 0.046 | 0.00355 | 1.871 |
| NW Eifuku | 38 | 23 | 19 | 0.899 ± 0.034 | 0.00472 | 2.615 |
| NW Rota | 332 | 63 | 103 | 0.899 ± 0.0103 | 0.00492 | 2.398 |
| Seamount X | 9 | 12 | 8 | 0.972 ± 0.064 | 0.00502 | 2.833 |
| All populations | 455 | 71 | 133 | 0.893 ± 0.0097 | 0.00487 | 2.369 |

**Table S3.** Genetic diversity based on partial COI sequences of *Rimicaris cambonae* and *Rimicaris loihi* for each sampling sites. **N:** number of sequenced individuals for each population; **S:** number of variable sites; **h:** haplotype number; **Hd:** haplotype diversity; **π:** nucleotide diversity; **k:** number of nucleotide differences.

| | N | S | h | Hd | π | k |
|---|---|---|---|---|---|---|
| ***Rimicaris loihi* from NW Rota** | | | | | | |
| 2004 | 62 | 29 | 27 | 0.913 ± 0.020 | 0.00478 | 2.646 |
| 2006 | 51 | 27 | 26 | 0.866 ± 0.036 | 0.00336 | 1.716 |
| 2009 | 48 | 20 | 19 | 0.877 ± 0.028 | 0.00467 | 2.272 |
| 2010 | 89 | 38 | 40 | 0.923 ± 0.016 | 0.00554 | 2.699 |
| 2014 | 82 | 31 | 35 | 0.864 ± 0.028 | 0.00325 | 1.770 |
| All NW Rota | 332 | 63 | 103 | 0.8986 ± 0.0103 | 0.00492 | 2.398 |

**Table S4.** Genetic diversity based on partial COI sequences of *Rimicaris loihi* from NW Rota for each sampling year at that vent field. **N:** number of sequenced individuals for each population; **S:** number of variable sites; **h:** haplotype number; **Hd:** haplotype diversity; **π:** nucleotide diversity; **k:** number of nucleotide differences.

| | *R. loihi:* Nikko | *R. loihi:* Suiyo | *R. loihi:* NW Eifuku | *R. cambonae:* NW Eifuku |
|---|---|---|---|---|
| *R. loihi:* Nikko | | 1,992,235 | 2,068,788 | 1,899,776 |
| *R. loihi:* Suiyo | 0 | | 1,825,580 | 1,645,704 |
| *R. loihi:* NW Eifuku | 0.075 | 0.013 | | 2,414,313 |
| *R. cambonae:* NW Eifuku | 0.175 | 0.103 | 0.121 | |

**Table S5.** Global estimates for weighted (ratio between alpha and alpha+beta) pairwise $F_{ST}$ calculated with realSFS for *R. loihi* and *R. cambonae* populations.

| | *R. loihi:* Nikko | *R. loihi:* Suiyo | *R. loihi:* NW Eifuku |
|---|---|---|---|
| *R. loihi:* Nikko | | | |
| *R. loihi:* Suiyo | 0.019 (0.015-0.022) | | |
| *R. loihi:* NW Eifuku | 0.061 (0.058-0.063) | 0.031 (0.029-0.034) | |
| *R. cambonae:* NW Eifuku | 0.163 (0.161-0.167) | 0.136 (0.133-0.138) | 0.109 (0.107-0.111) |

**Table S6.** Global estimates for pairwise $F_{ST}$ values calculated with StAMPP including upper and lower bounds in parentheses for *R. loih*i and *R. cambonae* populations. All estimates had significant p-values (p=0).

| Model | Log-L | AIC | theta | $Nu_{ca}$ | $Nu_{lo}$ | $Nu_{ca}b$ | $Nu_{lo}b$ | m | m12 | m21 | T/T1 | T2 | s | f |
|---|---|---|---|---|---|---|---|---|---|---|---|---|---|---|
| no_mig | -76810 | 153626 | 50864 | 0.06 | 0.10 | | | | | | 0.02 | | | |
| asym_mig | -77558 | 155126 | 50366 | 0.08 | 0.12 | | | | 0.92 | 0.02 | 1.5 | | | |
| asym_mig_size | -78715 | 157445 | 47437 | 6.67 | 0.24 | 0.05 | 0.72 | | 0.67 | 0.28 | 0.04 | 0.01 | | |
| sec_contact_asym_mig_size | -78766 | 157548 | 40248 | 3.85 | 2.91 | 0.27 | 0.21 | | 1.31 | 9.60 | 2.48 | 0.43 | | |
| anc_sym_mig_size | -80325 | 160665 | 45345 | 0.17 | 0.68 | 0.78 | 0.19 | 1.97 | | | 0.05 | 0.02 | | |
| vic_sec_contact_asym_mig | -80642 | 161295 | 18334 | | | | | | 1.13 | 4.18 | 9.83 | 0.74 | 0.47 | |
| vic_two_epoch_admix | -81526 | 163059 | 41985 | | | | | | | | 0.33 | 0.05 | 0.37 | 0.67 |
| vic_anc_asym_mig | -82635 | 165279 | 43194 | | | | | | 0.02 | 9.99 | 0.22 | 0.05 | 0.49 | |
| sec_contact_sym_mig | -82781 | 165572 | 16462 | 0.49 | 0.81 | | | 1.89 | | | 7.55 | 1.12 | | |
| sym_mig_size | -82984 | 165982 | 82208 | 0.54 | 11.19 | 0.10 | 0.16 | 9.22 | | | 0.62 | 0.34 | | |
| anc_sym_mig | -83570 | 167149 | 11725 | 1.48 | 1.97 | | | 5.19 | | | 26.35 | 0.31 | | |
| sec_contact_sym_mig_size | -83793 | 167600 | 66300 | 0.60 | 1.78 | 0.16 | 0.24 | 6.01 | | | 0.13 | 0.34 | | |
| founder_nomig_admix_two_epoch | -84095 | 168199 | 35204 | | 0.11 | | | | | | 1.69 | 0.03 | 0.03 | 0.85 |
| founder_nomig | -84453 | 168913 | 41481 | | 0.39 | | | | | | 0.10 | | 0.50 | |
| founder_sym | -84549 | 169106 | 41698 | | 0.36 | | | 0.03 | | | 0.10 | | 0.50 | |
| vic_no_mig | -84626 | 169257 | 40812 | | | | | | | | 0.10 | | 0.50 | |
| vic_no_mig_admix_early | -84629 | 169264 | 40810 | | | | | | | | 0.10 | | 0.50 | 0.97 |
| vic_anc_sym_mig | -84631 | 169269 | 40818 | | | | | 0.13 | | | 0.01 | 0.09 | 0.50 | |
| vic_no_mig_admix_late | -84856 | 169718 | 40804 | | | | | | | | 0.11 | | 0.50 | 0.01 |
| founder_nomig_admix_late | -85121 | 170250 | 42100 | | 0.31 | | | | | | 0.09 | | 0.50 | 0.03 |
| founder_asym | -85162 | 170334 | 42053 | | 0.32 | | | | 0.31 | 0.32 | 0.09 | | 0.50 | |
| vic_sec_contact_sym_mig | -85766 | 171541 | 25601 | | | | | 2.59 | | | 2.99 | 0.52 | 0.49 | |
| IM | -86542 | 173096 | 88627 | 0.17 | 0.18 | | | 3.12 | 8.15 | | 0.65 | | | 0.17 |
| sec_contact_asym_mig | -86720 | 173451 | 20382 | 0.24 | 1.24 | | | | 4.36 | 0.37 | 29.76 | 2.52 | | |
| founder_nomig_admix_early | -86768 | 173545 | 38931 | | 0.92 | | | | | | 0.12 | | 0.47 | 0.65 |
| anc_asym_mig | -87803 | 175617 | 8810 | 2.86 | 2.01 | | | | 0.39 | 1.39 | 23.35 | 0.23 | | |
| sym_mig | -90329 | 180665 | 19134 | 0.83 | 1.19 | | | 1.22 | | | 25.85 | | | |
| no_mig_size | -95841 | 191694 | 31482 | 6.20 | 5.42 | 0.58 | 0.56 | | | | 0.19 | 0.12 | | |
| anc_asym_mig_size | -96948 | 193912 | 32264 | 8.95 | 0.95 | 0.38 | 2.04 | 0.76 | 0.14 | | 0.17 | 0.07 | | |

**Table S7.** Unscaled demographic estimates and AIC for all models tested with dadi_pipeline for sympatric populations of *R. loihi* and *R. cambonae* from Eifuku seamount including: theta, mutation parameter for the ancestral population; nu, size of population after split; m, symmetrical migration; m12, migration from *R. loihi* into *R. cambonae*; m21, migration from *R. cambonae* into *R. loihi*, T1, time in the past of the split (2*Na generations); T2, time of population size change; s, fraction of Nref that goes to *R. loihi*, *R. loihi* size =s, *R. cambonae* size = 1-s; f, fraction of updated *R. loihi* to be derived from *R. cambonae*. Blank entries are parameters not estimated by model.

| Model | Log-L | AIC | theta | $Nu_{ca}$ | $Nu_{lo}$ | $Nu_{ca}$b | $Nu_{lo}$b | m | m12 | m21 | T/T1 | T2 |
|---|---|---|---|---|---|---|---|---|---|---|---|---|
| no_mig_size | -72098 | 144207 | 43313 | 0.69 | 0.06 | 0.13 | 22.76 | | | | 0.02 | 0.04 |
| sec_contact_asym_migsiz | -72888 | 145792 | 40012 | 1.38 | 0.24 | 0.12 | 3.32 | | 0.30 | 0.63 | 0.08 | 0.03 |
| no_mig | -73517 | 147041 | 43332 | 0.20 | 0.21 | | | | | | 0.06 | |
| anc_asym_mig | -73559 | 147130 | 43398 | 0.20 | 0.21 | | | | 0.66 | 3.57 | 0.02 | 0.05 |
| anc_sym_mig | -73829 | 147667 | 41172 | 0.26 | 0.27 | | | 0.97 | | | 0.02 | 0.06 |
| sym_mig | -74133 | 148274 | 42234 | 0.24 | 0.25 | 0.36 | 0.08 | | | | | |
| anc_sym_mig_size | -74421 | 148857 | 10563 | 2.12 | 1.83 | 0.66 | 1.01 | 1.17 | | | 9.31 | 0.15 |
| sec_contact_asym_mig | -74842 | 149696 | 40130 | 0.30 | 0.33 | | | | 0.54 | 0.05 | 0.01 | 0.09 |
| vic_two_epoch_admix | -75937 | 151882 | 38013 | | | | | | | | 0.07 | 0.15 |
| founder_nomig | -75996 | 151999 | 36493 | | 0.38 | | | | | | 0.14 | |
| vic_no_mig | -76106 | 152216 | 35723 | | | | | | | | 0.15 | |
| vic_no_mig_admix_ea | -76108 | 152223 | 35725 | | | | | | | | 0.15 | |
| vic_anc_sym_mig | -76112 | 152232 | 35716 | | | | | 5.65 | | | 0.02 | 0.14 |
| vic_no_mig_admix_late | -76168 | 152341 | 35721 | | | | | | | | 0.15 | |
| vic_anc_asym_mig | -76173 | 152356 | 35725 | | | | | | 1.14 | 5.45 | 0.05 | 0.12 |
| founder_asym | -76205 | 152419 | 37013 | | 0.33 | | | | 0.05 | 0.70 | 0.15 | |
| founder_sym | -76418 | 152843 | 36145 | | 0.43 | | | 0.16 | | | 0.15 | |
| founder_nomig_admix_early | -76776 | 153559 | 34773 | | 0.71 | | | | | | 0.16 | |
| founder_nomig_admix_late | -77270 | 154547 | 34137 | | 0.81 | | | | | | 0.17 | |
| sym_mig_size | -77564 | 155142 | 18684 | 1.53 | 0.75 | 0.31 | 0.61 | 1.13 | | | 7.33 | 0.09 |
| IM | -77972 | 155955 | 40518 | 0.15 | 0.36 | | | | 2.04 | 0.27 | 0.11 | |
| founder_nomig_admix_two_epoch | -79152 | 158314 | 53157 | | 0.26 | | | | | | 0.74 | 0.10 |
| vic_sec_contact_asym_mig | -81019 | 162048 | 26342 | | | | | | 0.43 | 3.10 | 8.21 | 1.43 |
| sec_contact_sym_mig | -83003 | 166016 | 13584 | 1.16 | 1.20 | | | 0.60 | | | 0.69 | 29.59 |
| vic_sec_contact_sym_mig | -83009 | 166026 | 32163 | | | | | 1.44 | | | 0.03 | 7.37 |
| asym_mig | -83022 | 166053 | 10817 | 1.51 | 1.47 | | | | 0.45 | 0.52 | 22.60 | |
| sec_contact_sym_mig_sz | -83039 | 166092 | 26898 | 1.72 | 9.80 | 0.59 | 0.61 | 1.25 | | | 0.02 | 23.94 |
| asym_mig_size | -83069 | 166154 | 27212 | 11.39 | 0.66 | 0.59 | 0.59 | | 1.18 | 1.15 | 0.69 | 1.71 |
| anc_asym_mig_size | -83736 | 167488 | 38443 | 0.69 | 0.24 | 9.78 | 1.59 | | 0.08 | 4.23 | 1.39 | 0.01 |

**Table S8.** Unscaled demographic estimates and AIC for all models tested with dadi_pipeline between allopatric populations of *R. loihi* from Nikko Seamount versus *R. cambonae* from Eifuku Seamount including: theta, mutation parameter for the ancestral population; nu, size of population after split; m, symmetrical migration; m12, migration from *R. loihi* into *R. cambonae*; m21, migration from *R. cambonae* into *R. loihi*, T1, time in the past of the split (2*Na generations); T2, time of population size change; s, fraction of Nref that goes to *R. loihi*, *R. loihi* size =s, *R. cambonae* size = 1-s; f, fraction of updated *R. loihi* to be derived from *R. cambonae*. Blank entries are parameters not estimated by model.

| Model | LogLn | AIC | theta | Nu1/ Nu1a | Nu2/ Nu2a | Nu1b | Nu2b | M | M12 | M21 | T/T1 | T2 | s | f |
|---|---|---|---|---|---|---|---|---|---|---|---|---|---|---|
| **Suiyo versus Nikko Seamounts** | | | | | | | | | | | | | | |
| anc_sym_mig_size | -65266 | 130546 | 39624 | 1.56 | 0.20 | 0.25 | 26.15 | 11.48 | | | 0.52 | 0.03 | | |
| vic_two_epoch_admix | -65601 | 131209 | 50137 | | | | | | | | 0.56 | 0.07 | 0.49 | 0.98 |
| founder_nomig | -65604 | 131214 | 33529 | | 3.78 | | | | | | 0.10 | | 0.22 | |
| founder_nomig_admix_ early | -65606 | 131220 | 33589 | | 3.40 | | | | | | 0.10 | | 0.23 | 0.12 |
| **Nikko versus Eifuku Seamounts** | | | | | | | | | | | | | | |
| vic_two_epoch_admix | -90068 | 180143 | 43944 | | | | | | | | 0.36 | 0.07 | 0.37 | 0.82 |
| no_mig_size | -90709 | 181429 | 41081 | 0.40 | 1.72 | 2.07 | 0.15 | | | | 0.09 | 0.02 | | |
| anc_sym_mig | -90734 | 181478 | 64643 | 0.23 | 0.26 | | | 27.43 | | | 0.47 | 0.04 | | |
| no_mig | -91050 | 182106 | 45339 | 0.28 | 0.29 | | | | | | 0.06 | | | |
| **Eifuku versus Suiyo Seamounts** | | | | | | | | | | | | | | |
| founder_nomig_admix_ two_epoch | -64740 | 129489 | 42155 | | 0.41 | | | | | | 0.25 | 0.05 | 0.49 | 0.98 |
| vic_two_epoch_admix | -64777 | 129562 | 41708 | | | | | | | | 0.24 | 0.05 | 0.49 | 0.98 |
| asym_mig | -65034 | 130077 | 79227 | 0.15 | 0.20 | | | | 29.51 | 1.59 | 0.30 | | | |
| sec_contact_asym_mig | -65292 | 130596 | 75502 | 0.16 | 0.26 | | | | 25.28 | 1.73 | 2.87 | 0.38 | | |

**Table S9.** The AIC and theta ($4N_{ref}ul$) estimates for the four most probable models and unscaled demographic parameters for dadi-pipeline analyses between each *R. loihi* population including: Nu, effective population sizes of (a) before the split and (b) after the split; m, migration; m12, migration from population 2 into population 1; m21, migration from population 1 into population 2; T, time of population split; T1, time in the past of the split (2*Na generations); T2, time of population size change; s, fraction of Nref that goes to population 1, size =s, Population 2 = 1-s; f, fraction of updated Population 1 to be derived from population 2. Blank entries are parameters not estimated by model.

| NCBI_ID | **Haplotype Number** | Loihi | Suiyo | Nikko | NW Eifuku | NW Rota | Seamount X | **Total** |
|---|---|---|---|---|---|---|---|---|
| ***Rimicaris loihi*** | | | | | | | | |
| DQ328819 | **Ol_Hap01** | 3 | 7 | 17 | 7 | 73 | 2 | **109** |
| DQ328820 | **Ol_Hap02** | 1 | 3 | 3 | 10 | 94 | | **112** |
| DQ328821 | **Ol_Hap03** | | | 1 | | 5 | | **6** |
| DQ328822 | **Ol_Hap04** | 1 | | | | | | **1** |
| DQ328823 | **Ol_Hap05** | 1 | | | | | | **1** |
| DQ328824 | **Ol_Hap06** | 1 | | | | 1 | 1 | **3** |
| DQ328825 | **Ol_Hap07** | | | 1 | | 1 | 1 | **3** |
| DQ328826 | **Ol_Hap08** | | | 1 | | 1 | 1 | **3** |
| DQ328827 | **Ol_Hap09** | | | | | 1 | 1 | **2** |
| DQ328828 | **Ol_Hap10** | | | | | 1 | | **1** |
| DQ328829 | **Ol_Hap11** | | | 1 | | 1 | | **2** |
| DQ328830 | **Ol_Hap12** | | | | | 3 | 1 | **4** |
| DQ328831 | **Ol_Hap13** | | | | | | | **1** |
| DQ328832 | **Ol_Hap14** | | 1 | | | 1 | | **2** |
| DQ328833 | **Ol_Hap15** | | 1 | | 1 | 4 | 1 | **7** |
| DQ328834 | **Ol_Hap16** | | | | 1 | | 1 | **2** |
| DQ328835 | **Ol_Hap17** | | | | 1 | | | **1** |
| DQ328836 | **Ol_Hap18** | | 1 | | 2 | 2 | | **5** |
| DQ328837 | **Ol_Hap19** | | | | 1 | | | **1** |
| PQ643539 | **Ol_Hap20** | | | | 1 | 21 | | **22** |
| PQ643540 | **Ol_Hap21** | | | | | 16 | | **16** |
| PQ643541 | **Ol_Hap22** | | | | 3 | 3 | | **6** |
| PQ643542 | **Ol_Hap23** | | | | 1 | 3 | | **4** |
| PQ643543 | **Ol_Hap24** | | 1 | | | 3 | | **4** |
| PQ643544 | **Ol_Hap25** | | | | | 4 | | **4** |
| PQ643545 | **Ol_Hap26** | | | | | 4 | | **4** |
| PQ643546 | **Ol_Hap27** | | | 1 | 2 | 1 | | **4** |
| PQ643547 | **Ol_Hap28** | | | | | 3 | | **3** |
| PQ643548 | **Ol_Hap29** | | | | | 3 | | **3** |
| PQ643549 | **Ol_Hap30** | | | | | 3 | | **3** |
| PQ643550 | **Ol_Hap31** | | | | | 2 | | **2** |

| | | | | | | | | |
|---|---|---|---|---|---|---|---|---|
| PQ643551 | **Ol_Hap32** | | | | | 2 | | **2** |
| PQ643552 | **Ol_Hap33** | | | 1 | | 1 | | **2** |
| PQ643553 | **Ol_Hap34** | | 1 | | | 1 | | **2** |
| PQ643554 | **Ol_Hap35** | | | | | 2 | | **2** |
| PQ643555 | **Ol_Hap36** | | 1 | | | 1 | | **2** |
| PQ643556 | **Ol_Hap37** | | | | | 2 | | **2** |
| PQ643557 | **Ol_Hap38** | | | | | 2 | | **2** |
| PQ643558 | **Ol_Hap39** | | | | | 2 | | **2** |
| PQ643559 | **Ol_Hap40** | | | | | 2 | | **2** |
| PQ643560 | **Ol_Hap41** | | | | | 2 | | **2** |
| PQ643561 | **Ol_Hap42** | | | | | 2 | | **2** |
| PQ643562 | **Ol_Hap43** | | | | | 1 | | **1** |
| PQ643563 | **Ol_Hap44** | | | | | 1 | | **1** |
| PQ643564 | **Ol_Hap45** | | | | | 1 | | **1** |
| PQ643565 | **Ol_Hap46** | | | | | 1 | | **1** |
| PQ643566 | **Ol_Hap47** | | | | | 1 | | **1** |
| PQ643567 | **Ol_Hap48** | | | | | 1 | | **1** |
| PQ643568 | **Ol_Hap49** | | | | | 1 | | **1** |
| PQ643569 | **Ol_Hap50** | | | | | 1 | | **1** |
| PQ643570 | **Ol_Hap51** | | | | | 1 | | **1** |
| PQ643571 | **Ol_Hap52** | | | | | 1 | | **1** |
| PQ643572 | **Ol_Hap53** | | | | | 1 | | **1** |
| PQ643573 | **Ol_Hap54** | | | | | 1 | | **1** |
| PQ643574 | **Ol_Hap55** | | | | | 1 | | **1** |
| PQ643575 | **Ol_Hap56** | | | 1 | | | | **1** |
| PQ643576 | **Ol_Hap57** | | | 1 | | | | **1** |
| PQ643577 | **Ol_Hap58** | | | 1 | | | | **1** |
| PQ643578 | **Ol_Hap59** | | | 1 | | | | **1** |
| PQ643579 | **Ol_Hap60** | | | 1 | | | | **1** |
| PQ643580 | **Ol_Hap61** | | | 1 | | | | **1** |
| PQ643581 | **Ol_Hap62** | | | 1 | | | | **1** |
| PQ643582 | **Ol_Hap63** | | | 1 | | | | **1** |
| PQ643583 | **Ol_Hap64** | | | 1 | | | | **1** |
| PQ643584 | **Ol_Hap65** | | | | 1 | | | **1** |

| | | | | | | | | |
|---|---|---|---|---|---|---|---|---|
| PQ643585 | **Ol_Hap66** | | | | 1 | | | **1** |
| PQ643586 | **Ol_Hap67** | | | | 1 | | | **1** |
| PQ643587 | **Ol_Hap68** | | | | 1 | | | **1** |
| PQ643588 | **Ol_Hap69** | | | | 1 | | | **1** |
| PQ643589 | **Ol_Hap70** | | | | | 1 | | **1** |
| PQ643590 | **Ol_Hap71** | | | | | 1 | | **1** |
| PQ643591 | **Ol_Hap72** | | | | | 1 | | **1** |
| PQ643592 | **Ol_Hap73** | | | | | 1 | | **1** |
| PQ643593 | **Ol_Hap74** | | | | | 1 | | **1** |
| PQ643594 | **Ol_Hap75** | | | | | 1 | | **1** |
| PQ643595 | **Ol_Hap76** | | | | | 1 | | **1** |
| PQ643596 | **Ol_Hap77** | | | | | 1 | | **1** |
| PQ643597 | **Ol_Hap78** | | | | | 1 | | **1** |
| PQ643598 | **Ol_Hap79** | | | | | 1 | | **1** |
| PQ643599 | **Ol_Hap80** | | | | | 1 | | **1** |
| PQ643600 | **Ol_Hap81** | | | | | 1 | | **1** |
| PQ643601 | **Ol_Hap82** | | | | | 1 | | **1** |
| PQ643602 | **Ol_Hap83** | | | | | 1 | | **1** |
| PQ643603 | **Ol_Hap84** | | | | | 1 | | **1** |
| PQ643604 | **Ol_Hap85** | | | | | 1 | | **1** |
| PQ643605 | **Ol_Hap86** | | | | | 1 | | **1** |
| PQ643606 | **Ol_Hap87** | | | | | 1 | | **1** |
| PQ643607 | **Ol_Hap88** | | | | | 1 | | **1** |
| PQ643608 | **Ol_Hap89** | | | | | 1 | | **1** |
| PQ643609 | **Ol_Hap90** | | | | | 1 | | **1** |
| PQ643610 | **Ol_Hap91** | | | | | 1 | | **1** |
| PQ643611 | **Ol_Hap92** | | | | | 1 | | **1** |
| PQ643612 | **Ol_Hap93** | | | | | 1 | | **1** |
| PQ643613 | **Ol_Hap94** | | | | | 1 | | **1** |
| PQ643614 | **Ol_Hap95** | | | | | 1 | | **1** |
| PQ643615 | **Ol_Hap96** | | | | | 1 | | **1** |
| PQ643616 | **Ol_Hap97** | | | | | 1 | | **1** |
| PQ643617 | **Ol_Hap98** | | | | | 1 | | **1** |
| PQ643618 | **Ol_Hap99** | | | | | 1 | | **1** |

| | | | | | | | | |
|---|---|---|---|---|---|---|---|---|
| PQ643619 | **Ol_Hap100** | | | | | 1 | | **1** |
| PQ643620 | **Ol_Hap101** | | | | | 1 | | **1** |
| PQ643621 | **Ol_Hap102** | | | | | 1 | | **1** |
| PQ643622 | **Ol_Hap103** | | | | | 1 | | **1** |
| PQ643623 | **Ol_Hap104** | | | | | 1 | | **1** |
| PQ643624 | **Ol_Hap105** | | | | | 1 | | **1** |
| PQ643625 | **Ol_Hap106** | | | | | 1 | | **1** |
| PQ643626 | **Ol_Hap107** | | | | | 1 | | **1** |
| PQ643627 | **Ol_Hap108** | | | | | 1 | | **1** |
| PQ643628 | **Ol_Hap109** | | | | | 1 | | **1** |
| PQ643629 | **Ol_Hap110** | | | | | 1 | | **1** |
| PQ643630 | **Ol_Hap111** | | | | | 1 | | **1** |
| PQ643631 | **Ol_Hap112** | | | | | 1 | | **1** |
| PQ643632 | **Ol_Hap113** | | | | | 1 | | **1** |
| PQ643633 | **Ol_Hap114** | | | | | 1 | | **1** |
| PQ643634 | **Ol_Hap115** | | | | | 1 | | **1** |
| PQ643635 | **Ol_Hap116** | | | | | 1 | | **1** |
| PQ643636 | **Ol_Hap117** | | | | | 1 | | **1** |
| PQ643637 | **Ol_Hap118** | | | | | 1 | | **1** |
| PQ643638 | **Ol_Hap119** | | | | | 1 | | **1** |
| PQ643639 | **Ol_Hap120** | | | | | 1 | | **1** |
| PQ643640 | **Ol_Hap121** | | | | | 1 | | **1** |
| PQ643641 | **Ol_Hap122** | | | | | 1 | | **1** |
| PQ643642 | **Ol_Hap123** | | | | | 1 | | **1** |
| PQ643643 | **Ol_Hap124** | | | | 1 | | | **1** |
| PQ643644 | **Ol_Hap125** | | | | 1 | | | **1** |
| PQ643645 | **Ol_Hap126** | | | | 1 | | | **1** |
| PQ643646 | **Ol_Hap127** | | 1 | | | | | **1** |
| PQ643647 | **Ol_Hap128** | | 1 | | | | | **1** |
| PQ643648 | **Ol_Hap129** | | 1 | | | | | **1** |
| PQ643649 | **Ol_Hap130** | | 1 | | | | | **1** |
| PQ643650 | **Ol_Hap131** | | 1 | | | | | **1** |
| PQ643651 | **Ol_Hap132** | | 1 | | | | | **1** |
| PQ643652 | **Ol_Hap133** | | 1 | | | | | **1** |

| | | | | | | | | |
|---|---|---|---|---|---|---|---|---|
| PQ643653 | **Ol_Hap134** | | 1 | | | | | **1** |

| NCBI_ID | **Haplotype Number** | NW Eifuku | NW Rota | Seamount X | West Mata | **Total** |
|---|---|---|---|---|---|---|
| ***Rimicaris cambonae*** | | | | | | |
| PQ643496 | **Oc_Hap01** | 98 | | 1 | 1 | **100** |
| PQ643497 | **Oc_Hap02** | 9 | | | | **9** |
| PQ643498 | **Oc_Hap03** | 4 | | | 4 | **8** |
| PQ643499 | **Oc_Hap04** | 3 | | | | **3** |
| PQ643500 | **Oc_Hap05** | 2 | | | | **2** |
| PQ643501 | **Oc_Hap06** | | 2 | | | **2** |
| PQ643502 | **Oc_Hap07** | 1 | | | | **1** |
| PQ643503 | **Oc_Hap08** | 1 | | | | **1** |
| PQ643504 | **Oc_Hap09** | 1 | | | | **1** |
| PQ643505 | **Oc_Hap10** | 1 | | | | **1** |
| PQ643506 | **Oc_Hap11** | 1 | | | | **1** |
| PQ643507 | **Oc_Hap12** | 1 | | | | **1** |
| PQ643508 | **Oc_Hap13** | 1 | | | | **1** |
| PQ643509 | **Oc_Hap14** | 1 | | | | **1** |
| PQ643510 | **Oc_Hap15** | 1 | | | | **1** |
| PQ643511 | **Oc_Hap16** | | 1 | | | **1** |
| PQ643512 | **Oc_Hap17** | | | | 1 | **1** |
| PQ643513 | **Oc_Hap18** | | | | 1 | **1** |
| PQ643514 | **Oc_Hap19** | | | | 1 | **1** |
| PQ643515 | **Oc_Hap20** | | | | 1 | **1** |
| PQ643516 | **Oc_Hap21** | | | | 1 | **1** |
| PQ643517 | **Oc_Hap22** | | | | 1 | **1** |
| PQ643518 | **Oc_Hap23** | | | | 1 | **1** |
| PQ643519 | **Oc_Hap24** | | | | 1 | **1** |

**Table S10.** haplotype number for partial COI sequences with corresponding NCBI ID for *Rimicaris loihi* (DQ328819 - DQ328837 and PQ643539 - PQ643653) and *Rimicaris cambonae* (PQ643496 - PQ643519).
